# Comparative mapping of crawling-cell morphodynamics in deep learning-based feature space

**DOI:** 10.1101/2020.09.06.285411

**Authors:** Daisuke Imoto, Nen Saito, Akihiko Nakajima, Gen Honda, Motohiko Ishida, Toyoko Sugita, Sayaka Ishihara, Koko Katagiri, Chika Okimura, Yoshiaki Iwadate, Satoshi Sawai

**Affiliations:** Department of Basic Science, Graduate School of Arts and Sciences, University of Tokyo, Meguro-ku, Tokyo 153-8902, Japan; Universal Biological Institute, University of Tokyo, Bunkyo-ku, Tokyo 113-0033, Japan; Research Center for Complex Systems Biology, Graduate School of Arts and Sciences, University of Tokyo, Meguro-ku, Tokyo 153-8902, Japan; Department of Biology, Graduate School of Science, University of Tokyo, Bunkyo-ku, Tokyo 1130033, Japan; School of Science, Kitasato University, Kitasato, Sagamihara, Kanagawa 252-0344, Japan; Faculty of Science, Yamaguchi University, Yamaguchi 753-8512, Japan; Exploratory Research Center on Life and Living Systems, National Institutes of Natural Sciences, Okazaki, Aichi 444-8787, Japan

## Abstract

Navigation of fast migrating cells such as amoeba *Dictyostelium* and immune cells are tightly associated with their morphologies that range from steady polarized forms that support high directionality to those more complex and variable when making frequent turns. Model simulations are essential for quantitative understanding of these features and their origins, however systematic comparisons with real data are underdeveloped. Here, by employing deep-learning-based feature extraction combined with phase-field modeling framework, we show that a low dimensional feature space for 2D migrating cell morphologies obtained from the shape stereotype of keratocytes, *Dictyostelium* and neutrophils can be fully mapped by interlinked signaling network of cell-polarization and protrusion dynamics. Our analysis links the data-driven shape analysis to the underlying causalities by identifying key parameters critical for migratory morphologies both normal and aberrant under genetic and pharmacological perturbations. The results underscore the importance of deciphering self-organizing states and their interplay when characterizing morphological phenotypes.

## Introduction

Cell migration is a fundamental cellular process that underlies embryonic development, wound healing, immunological surveillance and cancer metastasis. In particular, fast migrating cells such as *Dictyostelium* and migrating immune cells are versatile in their patterns of movement that ranges from random exploratory movements with frequent turns to more persistent migration in a straight path. *Dictyostelium* cells exhibit random migration (*1*) as a phagocyte, in addition to more persistent migration as they aggregate under starvation to form a fruiting body. Exploratory interstitial migration in leukocytes (*2, 3*) underlies antigen search and immune surveillance (*4*), and some are also known to move in a straight line (*5*). Frequency of cell turning and their angles is dictated by when and where branched networks of F-actin that drives formation of lateral protrusions called pseudopods occur. In *Dictyostelium*, pseudopods appear arm-like, and their formation and splitting randomizes cell orientation (*6*). Selective maintenance of pseudopods thus provides directional bias in the shallow attractant gradients (*7*). Similar F-actin enriched projections in immune cells vary in their appearance from those that are finger-like in DC cells to those more lamellar in neutrophils, however their role in directional choice appears to be conserved (*8, 9*). On the other hand, ability to move in a straight and persistent manner requires cell polarity which refers to a long-term state having a dominant leading edge enriched in branched F-actin meshwork and a trailing end with crosslinked actomyosin. In certain cells under geometrical confinement, buildup of hydrostatic pressure by contractility can rapidly switch the protrusion to a bleb which is devoid of F-actin (*10*). Besides such cases, F-actin driven leading edge protrusion and rear contraction are concomitant in the polarized cells. While the particular shape that cells take depends on the extracellular conditions such as cell-substrate adhesion and diffusible attractants, shapes with broken-symmetry emerge in the absence of extracellular asymmetries and thus their origins are cell-intrinsic by nature (*6*, *7*, *11*). The fact that movement of fast migrating cells depend highly on self-deformation contrasts highly to those of mesenchymal cells such as fibroblasts, which move at an order of magnitude slower speed and are strongly dictated by the asymmetries introduced by the adhesive foci.

The common and recurring shapes observed under highly divergent culture conditions and across evolutionary distant species and taxa (*12*) suggest generality of the self-deforming dynamics in fast migrating cells. Transmigrating neutrophils, genetically or pharmacologically perturbed *Dictyostelium* (*13*, *14*) and certain cancer cells (*15*) take a canoe-like polarized morphology similar to fish-keratocytes and some protozoan amoebae. Conversely, polarized neutrophils under certain genetic and pharmacological perturbations are known to exhibit increased number of pseudopods (*16*, *17*). These common and interconvertible morphologies suggest that they reflect basic self-organizing states of motile cells that can be quantified and compared with minimal reference to the details of the molecular underpinnings and the permissive extracellular conditions. Due to compounding levels of complexity, quantitative characterization of these canonical morphologies also requires one to leave aside fine-scale protrusions such as filopodia and endocytic cups and apply an appropriate coarse-grained description at the cellular-level. The aspect ratio of fish keratocytes was identified as the major variation in shape features by principle component analysis (PCA) (*18*). Variation in more complex features requires other non-trivial measures of characterization. Fourier and related spectral analysis allows one to extract the periodicity in the protrusion-retraction cycle as well as in their spatial ordering (*19*). Combined with PCA, Fourier description of *Dictyostelium* cell shape has shown that the morphologies observed at various steepness of a chemo-attractant gradient can be characterized in a two-dimensional feature space that represents differences in degree of elongation, splitting and polarization (*20*). Zernike polynomials in combination with PCA has been used to classify invasive cancer morphologies in two-dimensional feature space (*21*). Besides Fourier-based analysis, methods such as tracking of local curvature (*22*) and pseudopods at the cell edge (*7*, *23*) have been employed to characterize spatio-temporal dynamics of membrane protrusions. In RNAi screen of *Drosophila* culture cells, a large body of hand-picked morphology features has been employed to train a classifier by shallow neural networks(*24*) and Support Vector Machine (*25*). These studies indicate that the states of physically realizable morphologies are confined to a relatively low-dimensional feature space (*25*). The downside of data-driven approaches, however, are that the analysis often remains in a black-box making it difficult for one to understand data with reference to the underlying causalities.

A great challenge remains as to how one can quantitatively relate the characterized shapes to the underlying dynamics and vice versa (*12*, *26*, *27*). For the most basic analysis, it is instructive to formulate a top-down model for isolated cells that is free of extracellular context (*12*), as behaviors under complex environments may later be deduced, given the repertoire of realizable dynamics, from spatially asymmetries and constraints in the key parameters. In the “graded radial extension model”, a polarized morphology similar to that observed in fish keratocytes and neutrophils is described without reference to the underlying mechanism by assuming that the plasma membrane extends radially and that its magnitude is spatially graded along the anterior-posterior axis (*28*). Such a steady and graded distribution is thought to result from reaction-diffusion based symmetry breaking in the activity of the polarity signals GTPases Rac and RhoA at the plasma membrane that specifies the state of F-actin at each given place and time. Resource limitation that prevents one state from dominating the other is expected when the sum of the inactive and active form of the small GTPases is approximately fixed in time (*29*). Bi-stable reaction-diffusion systems with the above constraint are known to support a protrusive membrane region (front) and contractile membrane region (rear) to co-exist in a spatially separate domains within a cell - a mathematical manifestation of a stable polarized cell shape (*29*). On the other hand, pseudopods are transient structures regulated by locally amplified formation of branched F-actin networks. In *Dictyostelium* this is governed by transient activation in Ras/Rap and PI3K (*30*), and in case of neutrophils, by Cdc42 and PI3K (*31, 32*). Because the localized protrusive dynamics occur under uniform conditions, they are thought to arise by noise amplification by excitable regulatory network (*23*, *30*, *32*, *33*). Cdc42 in neutrophils and Ras/Rap in *Dictyostelium* are also known to act positively to strengthen Rac and Rho and hence cell polarity (*16, 34–36*). Mathematical models of random turning and persistent migration thusfar have descried the outcome of either excitability-based (*14*, *37*) or polarity-based regulation (*38*), however realistic shapes resulting from the interplay of cell polarity and pseudopods have not been addressed not to mention the lack of systematic and quantitative morphology comparison between simulations and real data. In this work, to overcome these shortcomings, we first developed a framework that employs deep learning based classifier to obtain objective measures for shape comparison. To train neural networks, we chose the three well-studied fast migrating cells, *Dictyostelium*, neutrophils and keratocyte which have contrasting degree of cell polarity and pseudopods. The obtained feature space is then analyzed by formulating and computing a model of cell deformation that incorporates a conceptual signaling network based on the current knowledge of the pseudopod and cell polarity dynamics. Our analysis indicates that the proposed dynamics successfully map experimentally observed morphologies across the full range of the feature space and highlights key parameters that define morphologies under normal and aberrant conditions. The present approach provides a general and extendable framework to characterize varieties of other cell shapes in a data-driven manner which can then be interpreted and tested through hypothesis-driven modeling. Provided that there are large sets of simulated timeseries and real data for feature extraction, the ability to help infer the migratory dynamics from snapshot images should also have practical single-cell applications for cell identification.

## Results

### A feature space related to cell polarity and pseudopod dynamics can be obtained from classification of stereotype morphologies by deep convolutional neural networks

For systematic extraction of cell morphology features from microscopy data, a convolutional neural network (Fig. 1a, lower panel) was trained to classify shape of a cell based on whether it exhibited front- to-tail elongation with 1) a frequent or 2) occasional pseudopods or whether they took 3) a gliding form elongated in the lateral direction. To this end, we used snapshot images of well-studied cell types known for each of the above stereotype morphologies: *Dictyostelium* (aggregation-stage; ‘agg’), neutrophil-like HL-60 and fish keratocytes (Fig. 1a, upper panel; Movie S1). The choice of the reference data was based on the fact that they are well-studied systems and that each represented a different degree of pseudopod formation and cell polarity (*6*, *12*). Under our experimental conditions, *Dictyostelium* cells showed an elongated form in the anterior to posterior direction with locally appearing pseudopods. Fish keratocyte took a canoe-like shape characterized by its long axis orthogonal to the moving direction. HL-60 exhibited an intermediate form between the two where, compared to *Dictyostelium*, transient protrusions appear less frequently and the overall shape was more horizontally elongated but to a lesser extent than the keratocyte. Image masks of these isolated single cells (Table S1) were normalized in size and orientation (Fig. 1a; see Methods). Hyper-parameters for deep-learning were chosen for relative high-accuracy for various network structures (Materials and Methods). The extracted features were well trained as judged by the high validation accuracy; 97.9% and 89.7%, for the training and the validation data respectively (fig. S1a,b; the mean of the last 10 epochs). The three nodes ***F*** = (*F*_1_, *F*_2_, *F*_3_) that constituted the second to last layer of the network showed good representation of the three data classes: *Dictyoselium* (fig. S1c; high *F*_1_), HL-60 (fig. S1d; high *F*_2_) and keratocyte (Figure S1e; high *F*_3_), which can further be reduced to two by principal component analysis (PCA). The latency values of PC1, PC2 and PC3 were approximately 66.3%, 33.5% and 0.3%, respectively. PC1 = (−0.42,-0.35, 0.84)•***F*** and PC2 = (0.76,-0.64, 0.11)•***F.*** Figure 1b shows good separation of the three datasets in the PCA space. The keratocyte and *Dictyostelium* (agg) dataset were found in the high PC1 and high PC2 regions, respectively. The HL-60 dataset were mapped to a low PC1 low PC2 region.

**Figure 1.**
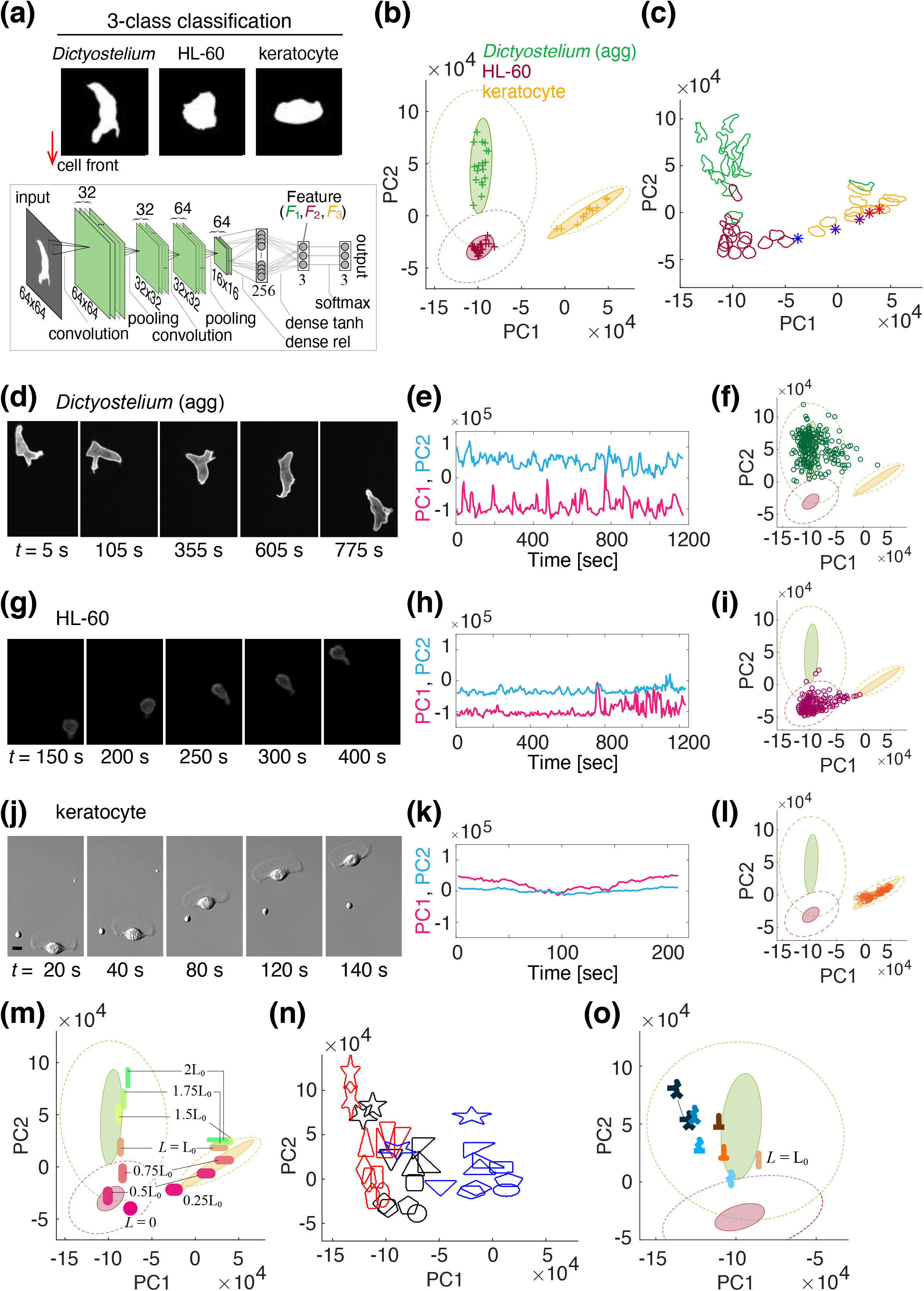
Stereotype migrating cell morphology can be classified in two-dimensional feature space. (a) Representative mask images (upper panels: *Dictyoselium* (agg; aggregation-stage), HL-60 and fish keratocyte) used for trainining a deep convolutional neural network (lower panel). Masks were normalized in area and aligned downwards in the migrating direction. The feature vector ***F*** = (*F*_1_, *F*_2_, *F*_3_) defined by the three nodes in the last layer was further reduced to PC1 and PC2 by PCA. (b) Mapping of trained data in PC1 and PC2 (dark green + (aggregation-stage *Dictyostelium*), dark red + (HL-60) and yellow + (keratocyte)). Each data point represents time-averaged scores from a time-series of a single cell. (c) Representative cell contours mapped to the feature space. Asterisks are the first principal shape variation of fish keratocyte (images taken from (*18*) (−2σ (blue) to +2σ from the mean (purple)). (d-l) Representative time-series and the corresponding feature scores for *Dictyostelium* (agg) (d-f), HL-60 (g-i) and fish keratocyte (j-l), respectively. (m-o) Mapping of skewed ellipsoidal shapes (the number indicates aspect ratio) (m), polygons (n) and a multi edge geometry (o) in the feature space. The circled regions in the background (b, f, i, m, o) are 95% confidence ellipses for the mean of all timeseries combined (dotted) and for the mean of individual cell (filled); green (*Dictyostelium*), dark red (HL-60), yellow (keratocyte).X

The feature metrics acquired above, by the very fact that they constitute a good classifier, should be useful to quantify similarity of cell morphology of one’s interest in reference to the trained data. The overall relationship between representative cell contours and the morphology feature is shown in Figure 1c. At first glance, higher PC1 appears to indicate more pronounced elongation in the lateral direction while higher PC2 indicates marked longitudinal elongation and protruding edges. These variations in the feature space not only reflected the average morphology differences between the three training classes but also shape changes in time. Figures 1d-l show representative single-cell timeseries from the validation dataset; i.e. a reserved dataset not used for training. *Dictyostelium* data showed large fluctuations in both the PC1 and PC2 direction (Fig. 1d-f), whereas the HL60 (Fig. 1g-i) and keratocyte data (Fig. 1j-l) exhibited more marked changes in the PC1 direction (see also fig. S1c-e for changes in ***F***). To clarify the relationship between the PC1-PC2 and the extent of elongation, we re-visited the principle component of the fish keratocyte shape variation reported earlier (‘shape mode 1’ in (*18*)). Low aspect-ratio shapes (*-2σ* in ‘shape mode 1’ (*18*)) were located near the HL-60 dataset, and high aspect-ratio shapes (2*σ* in ‘shape mode 1’ (*18*)) were located near the keratocyte dataset. Consequently, the shape along the ‘shape mode 1’ of fish keratocyte (*18*) constitutes a well-confined manifold in the PC1-PC2 space (Fig. 1c; asterisks) to which our fish keratocyte data were also mapped. Furthermore, ellipsoidal shapes with various aspect ratios indicate that an increase in the lateral elongation maps them on the same manifold as the keratocyte dataset (Fig. 1m, PC1 > −1; magenta to pink ellipsoids) while an increase the degree of head-to-tail elongation maps the ellipsoids to another manifold along the PC2 axis (Fig. 1m, green ellipsoids). Rotating these ellipsoids to intermediate angles map these shapes to intermediate PC1 values (fig. S1 f,g).

While the analysis of the stretched ellipsoids clearly demonstrated how the cell orientation and the degree of elongation are mapped to the feature space, the *Dictyostelium* (agg) dataset was markedly offset from these ellipsoids towards negative PC1 values. To further clarify the nature of this deviation, well-defined polygons were subjected to the same feature analysis (Materials and Methods). All 33 geometrical objects tested were mapped within the region of the PC1-PC2 space spanned by the microscopy data (Fig. 1n). Regular polygons and circles were mapped to a low PC1 - PC2 region and their vertically stretched counterparts were mapped to higher PC2 region (Fig. 1n, red). Polygons and ellipsoids that were stretched in the lateral direction marked high PC1 value (Fig. 1n, blue; figure S1h,i). Of particular note were the star objects which scored highest in the PC2 value and deviated markedly in the feature space from the ellipsoids and other objects when stretched (fig. S1i). Stars have higher PC2 compared to squares and triangles indicating that PC2 reflects pointed edges. Comparisons between upright and vertically flipped stars and triangles indicated that degree of pointedness towards the cell front also affects PC2 but in a complex way (Fig. 1n; see also fig. S1j). An analysis of more asymmetrical geometries with varying number of edges showed that they map to a domain in high PC2 with high variations towards negative PC1 values as in the *Dictyostelium* (agg) dataset; i.e. away from ellipsoidal shapes (Fig. 1o). Rotating the multi-edge forms with the small PC1 values (Fig. 1o; 0 to 30 degrees) bring about decrease in PC1 (fig. S1k). Further rotation increases PC1 (fig. S1k; 60 to 90 degrees) due to the shape now appearing more horizontally elongated overall. Generality of the results was confirmed in independent real-cell data by analyzing fully differentiated prespore cells of *Dictyostelium* which is elongated longitudinally and lacks pseudopods thus mapping identically to the elliposoids (fig. S2a, b). Likewise, effector T cells with their signature branching protrusions were broadly distributed along PC2 (fig. S2c, Th1) compared to markedly less polarized regulatory T cells (fig. S2d, Treg). These analysis indicate that our classifier was able to yield data-driven representation of complex signatures with respect to the cell orientation for both local pseudopodal protrusions and more global cell elongation.

### The ‘ideal cell’ model recapitulates a generalized morphological landscape constrained by the choice of the protrusion speed and the balance between the local protrusion and global polarity

Let us now introduce an ideal cell model (see Equations: Eqs. 1 and 2) that serves as a canonical shape generator of migrating cells that is strictly constrained by the transient protrusion dynamics and polarization (Supplementary Text). To describe the interfacial membrane mechanics, we employed the phase-field method with the addition of an active force *F*_prot_ = *a*_w_*W* (Eq. 1). The model also consists of spatio-temporal dynamics of variable *W* and variables *U*, *V* that define global cell polarity and local protrusions, respectively (Fig. 2a; Eq. 2). These variables are abstract representation of how the respective signaling molecules such as Rac, Rho for *W* and Ras, Cdc42, PI3K for *V* are regulated in space and time (Materials and Methods). *W* is bistable and takes either a low or a high state which signifies the retracting rear and the expanding front, respectively. The dynamics of *W* therefore supports a polarized shape with *W* being high at one end and low at the other end (Fig. 2b, *t* = 660s). The low state indicates that *W* has converted to the other form *W*\* while the integrated sum of *W* and *W*\* is fixed to *W*_tot_. In addition, there is an excitable network that describes conversion of *U* to *V* which is invoked by small signal fluctuations. The important assumption in the model is that these two core networks are coupled so that *V* promotes conversion of *W* from the low to the high state, and *W* catalyzes amplification of *V* (Fig. 2a). Because *V* amplifies noisy fluctuations and generates local protrusions through *W*, a leading edge defined by a region with high *W* is most likely to be perturbed and often split into two (Fig. 2b; *t* = 700). This is however transient, as *W* by itself works to maintain global unipolarity hence only one protrusion survives (Fig. 2b; *t* = 800s). In some cases, a new protrusion can also form away from the anterior and more towards the lateral side and still develop into a new dominant front (Fig. 2c). These features required full 3-variable equations (Eq. 2; Table S3) and were of particular importance in our comparative analysis. Insofar as our parameter search (Table S4 and S5), neither the dynamics of *U* and *V* (fig. S3a-d) nor that of *W* alone (fig. S3e-i) supported these bifurcating protrusions (Supplementary Text). Although our model encompasses the 1- and 2-variable limits which well describe morphologies outside of the training dataset such as oscillatory non-migratory shapes (fig. S3b,c) and non-bifurcating polarized cells (fig. S3f,g), we shall exclude these parameter regimes from the following analysis. Related oscillatory and fan-like morphologies have been addressed earlier (*39*–*41*).

**Figure 2.**
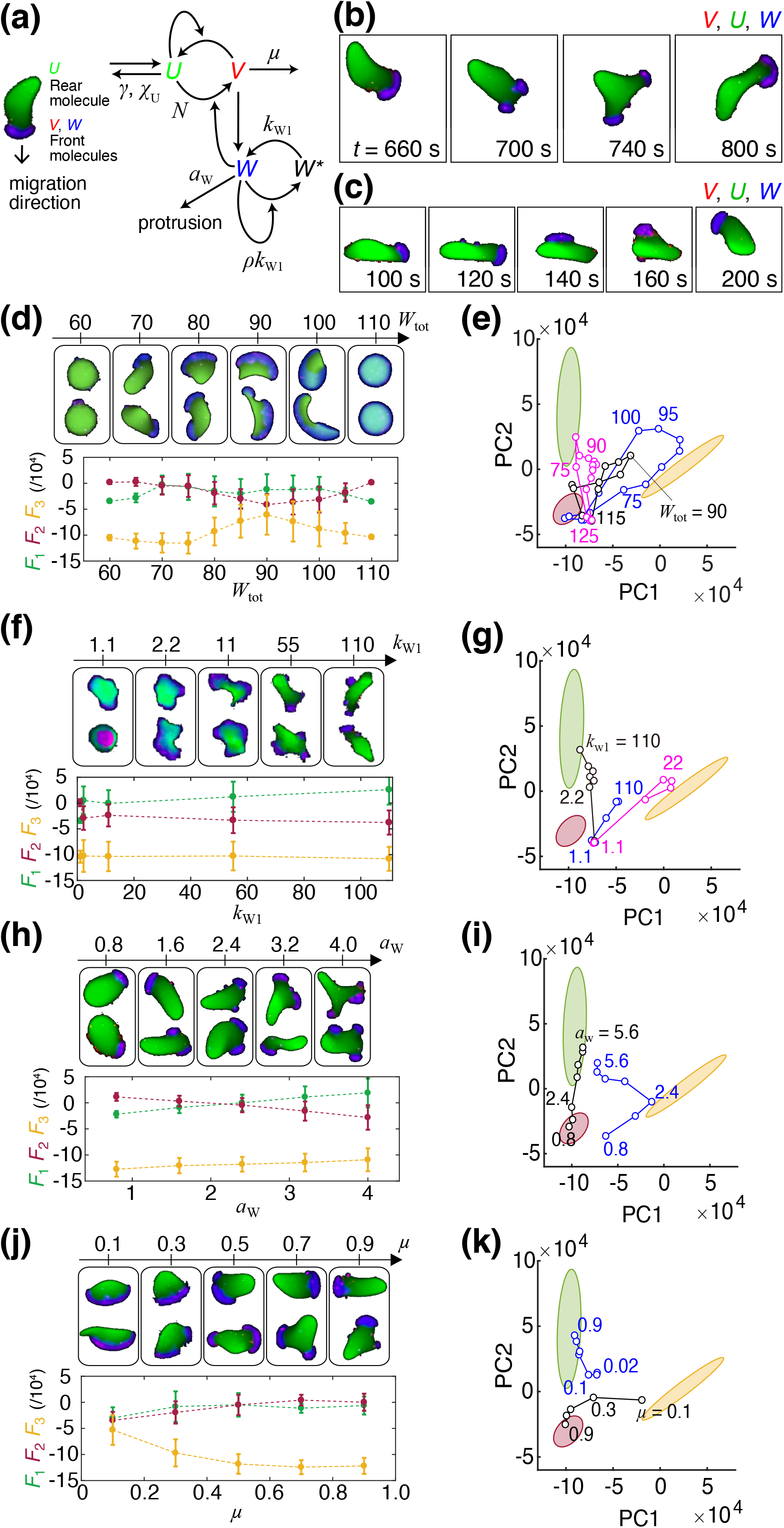
Degree of leading edge expansion and the balance between pseudopod formation and polarity persistence can account for the main morphology feature. (a) Schematics of the dynamical model. Self-amplifying and excitatory synthesis from *U* to *V* (*67*) induces protrusion factor *W* which can be prolonged by the bistable dynamics (Eq. 2). The reaction takes place in a region *ϕ* = 1 governed by interface physics according to the phase-field equation (Eq. 1). The membrane expands outward into an unoccupied region *ϕ* = 0 in the direction perpendicular to the border at a rate proportional to local *W*. Main parameters (*χ*_U_, *γ*, *μ*, *k*_W1_, *ρ*, *a*_W_) are denoted along the associated reaction steps. (b,c) Representative model behavior (overlay plots for *U*, *V*, *W*); front splitting (b) and pseudopod formation (c). (d-k) Parameter dependence of cell morphology (d,f,h,j upper panels), the average feature ***F*** from 2-4 independent simulation runs (d,f,h,j lower panels) and changes in the PC1-PC2 space; (d) *W*_tot_ (*χ*_U_ = 50, *k*_W1_ = 90, *μ* = 0.5, *ρ* = 5.5556, *γ* = 0.1, *a*_W_ = 2.4, *D*_W_ = 3). (e) Black same as (d), blue (*χ*_U_ = 50, *k*_W1_ = 100, *μ* = 0.1, *ρ* = 5, *γ* = 0.1, *a*_W_ =2.4, *D*_W_ = 3) and magenta (*χ*_U_=50, *k*_W1_=110, *μ*=0.5, *ρ*=4.5455, *γ*=0.1, *a*_W_ =4, *D*_W_=3). (f) *k*_W1_ (*χ*_U_ = 50, *μ* = 0.5, *ρ* = 4.5455, *γ* = 0.1, *a*_W_ = 5.6, *D*_W_ = 3, *W*_tot_ = 80). (g) Black same as (f), blue (*χ*_U_ = 50, *μ* = 0.5, *ρ* = 4.5455, *γ* = 0.1, *a*_W_ = 2.4, *D*_W_ = 3, *W*_tot_ = 90), and magenta (*χ*_U_ = 50, *μ* = 0.1, *ρ* = 4.5455, *γ* = 0.1, *a*_W_ = 2.4, *D*_W_ = 3, *W*_tot_ = 80). (h) *a*_W_ (*χ*_U_ = 30, *k*_W1_ = 110, *μ* = 0.5, *ρ* = 4.5455, *γ* = 0.1, *D*_W_ = 3, *W*_tot_ = 80). (i) Black same as (h), blue (*χ*_U_ = 50, *k*_W1_ = 100, *μ* = 0.1, *ρ* = 5, *γ* = 0.1, *D*_W_ = 3, *W*_tot_ = 80). (j) *μ* (*χ*_U_=50, *k*_W1_=100, *ρ*=5, *γ*=0.1, *a*_W_ =2.4, *D*_W_=3, *W*_tot_=80). (k) Black same as (j), the blue line is (*χ*_U_ = 50, *k*_W1_ = 110, *ρ* = 4.5455, *γ* = 0.1, *a*_W_ = 4.8, *D*_W_ = 3, *W*_tot_ = 80).

In total, 228 parameter conditions (Table S6) were selected by heuristic sampling where we performed grid search in the parameter subspace around hand-picked reference points (fig. S4; Material & Methods). The advantage of the ‘ideal cell’ model over the existing models focused on specific cells and conditions is that, it can describe a generalized morphological landscape incurred by the choice of the protrusion speed and the balance between the local protrusion and global polarity. As we shall describe later, this versatility allows us to resolve and characterize cells in different developmental stages and under perturbations. First, of particular interest was whether the cell was elongated in the lateral or anterior-posterior direction. We found *W*_tot_ has a large influence in the direction and the degree of overall cell elongation (Fig. 2d; see also Movie S2). For small *W*tot, the simulated morphology was near circular (Fig. 2d, *W*_tot_ = 50-70). As *W*_tot_ increased, a small region with *W* appeared, and cells became elongated (Fig. 2d, *W*_tot_ = 80, 90). A further increase in *W*_tot_, expanded the high *W* region (Fig. 2d, *W*_tot_ = 100) until it encompassed the entire perimeter (Fig. 2d, *W*_tot_ = 110). Accordingly, moderately high *W*_tot_ (*W*_tot_ = 80) supported relatively high *F*_1_ (indicating *Dictyostelium*-like shape) (Fig. 2d; bottom panel). In the PC1-PC2 space, small *W*_tot_ yielded low PC1 and low PC2, whereas at a moderately high *W*_tot_, PC2 took high values (Fig. 2e). At high *W*_tot_, PC1 and PC2 decreased. The overall dependency on *W*_tot_ were conserved when other parameters were varied (Fig. 2e; magenta and blue). The other important parameters that affected cell polarity was *ρ*; the conversion rate from W to W*. High *ρ* means that the non-zero roots of the cubic equation ***-ρW*^3^ + *ρW*W*^2^ - *W* = 0** is large thus supporting a larger domain with high *W*. Therefore, at low *ρ*, the leading edge is small and cells become elongated in the moving direction (fig. S5a; *ρ* = 4.55). As *ρ* is increased, leading edge became broader and *F*_3_ increased (fig. S5b;*ρ* = 5.56).

Occurrence of local protrusions depended largely on *k*_W1_ and *a*_W_. *k*_W1_ specifies the depth of the bistable well. For small *k*_W1_, the front-rear asymmetry was weak, and the overall cell shape was near circular (Fig. 2f; *k*_W1_ = 1.1). Intermediate *k*_W1_ exhibited mixed dynamics where local protrusions induced asymmetrical deformation however without persistent front-to-back polarity (Fig. 2f; *k*_W1_ = 2.2 – 55). Large *k*_W1_ elevates *W* which makes it less affected by the dynamics of *U* and *V*, and thus supports elongated shape with more marked polar asymmetry in *W* (Fig. 2f, *k*_W1_ = 110; Movie S3). Accordingly we obtained high *F*_1_ (Fig. 2f, lower panel), and PC2 (Fig. 2g). The appearance of local protrusions also depended strongly on the protrusion force *a*_W_ (Fig. 2h; Movie S4). For low *a*_W_ (Fig. 2h; *a*_W_ = 0.8), only small deformation was observed and the overall cell shape was near circular. At an intermediate value of *a*_W_ (Fig. 2h; *a*_W_ = 1.6 - 2.4), cells were more longitudinally elongated and the cell displacement was more directional. At high *a*_W_ (Fig. 2h; *a*_W_ = 3.2 – 4.0), multiple pseudopods appeared, and the cell orientation changed frequently (Movie S4). There was an increase and a decrease in *F*_1_ and *F*_2_ respectively (Fig. 2h, lower panel). The PC2 score increased accordingly (Fig. 2i).

Parameters that affected the pseudopod dynamics were *μ* and *γ* which define the downregulation rate of *V* and *U*, respectively. Low *μ* (Fig. 2j; *μ* = 0.1; Movie S5) elevates *V*, hence the concomitant increase in *W* supported a laterally elongated shape. The polarized shape was highly persistent as high *V* renders the patterning less prone to noise perturbation. On the other hand, at intermediate to high value of *μ*, cells became more elongated longitudinally and the polarity was less persistent (Fig. 2j; *μ* =0.5 - 0.9). Here, the high *W* domain was easily disrupted; fronts frequently split, and the cell orientation was altered (e.g., a Y-shaped front in Fig. 2j at *μ* = 0.7 and 0.9). Accordingly, *F*_3_ (Fig. 2j; bottom panel) and the PC1 score decreased at high *μ* (Fig.2k). Similarly, at low *γ*, pseudopods split frequently and new pseudopods were rare (fig. S5c, d; *γ* = 0.1) and the opposite was true for high *γ* (fig. S5c, d; *γ* = 0.5 or 0.7). Additionally, for splitting to occur, it was important that diffusion of *W* does not average out the local perturbations. Broad leading edge split at low *D*_W_, (fig. S5e; *D*_W_ = 0.6, 1.8 and 2.4), but was sustained at high *D*_W_ (fig. S5e; *D*_W_ = 3.6). These details only made subtle changes in our morphology feature (fig. S5e,f). The boundary flux *χ*_U_, was also important to restrict the *U-V* reaction at the edge (fig. S3b,c). Splitting of the front occurred more frequently at high *χ*_U_ (fig. S5g). Due to the temporal nature, the feature vector on average remained almost unchanged (fig. S5h).

### Morphology-based mapping of model parameters can help infer candidate dynamics

The distribution of the simulation data in the PC1-PC2 space were found to span a large region occupied by the training dataset (Fig. 3a, black circles) further vindicating the ability of the ideal cell model to describe the characteristic morphologies. Proximity of the time-averaged simulated morphologies (Fig. 3b) to the average of the three reference dataset was analyzed by computing the Euclidean distance in the feature space ***F*** = (*F*_1_, *F*_2_, *F*_3_). According to the reference data, the distance was designated as Score-D (*Dictyostelium*), Score-H (HL-60), Score-K (keratocytes), and ranked in the ascending order; i.e. a low score means high similarity (Table S7). The time averaged morphology feature in the PC1-PC2 space and the time-series of the top ranking simulations are shown in Fig. 3b (filled circles) and Figures 3c-k, respectively. Simulations with high Score-D on average exhibited morphology that closely resembled the aggregation-stage *Dictyostelium* with their elongated form in the anterior-posterior direction accompanied by a few pseudopods that frequently reoriented cell directionality (Fig. 3c; Movie S6). Similarly, simulations that ranked high for Score-H (Fig. 3f; Movie S7) exhibited fan-like cell shape that moved directionally with some occasional turning as observed in HL-60. For high Score-K simulations, cells had canoe-like shape with high directional persistence (Fig. 3i; Movie S8), however the similarity was not as high as the other two dataset. The top ranking parameter sets were found near the median of the reference dataset in PC1-PC2 space except for the keratocyte shape which deviated for the simulation in the PC2 direction. In our parameter search, we were unable to obtain simulation results that mapped closer to the average of the keratocyte data. The overall mapping of real cell data and model simulations were conserved when intermediate layer of the classifier was used to obtain the feature space (Supplementary Text; fig. S6; SI Table 9).

**Figure 3.**
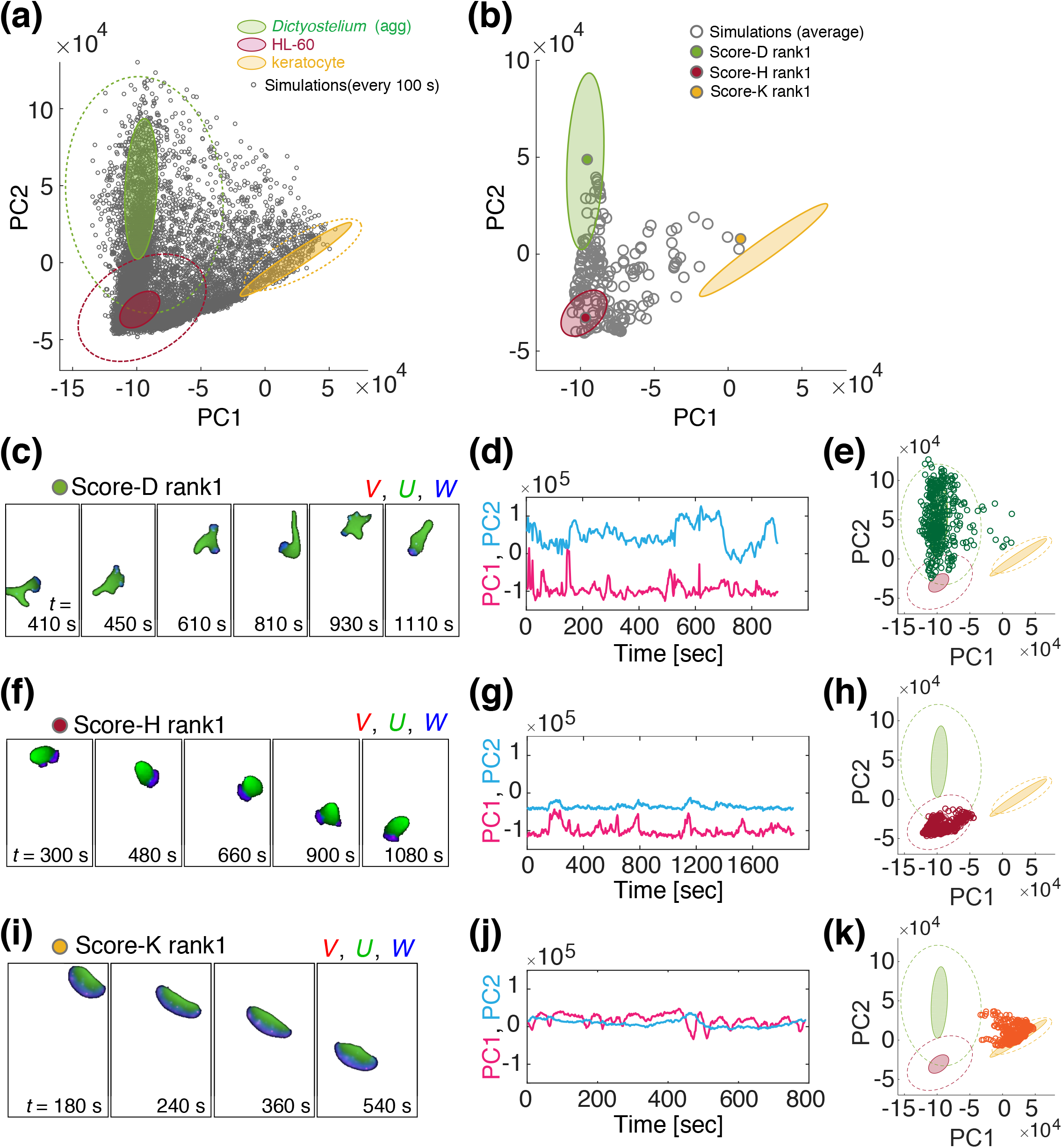
Mapping of morphology features between simulated and real cell data confirms critical parameters that define the morphology type. (a, b) Mapping of simulated cell morphology from the 228 parameter sets (Grey open circle: subsampled time series (a) and time averages for 2 to 4 independent runs (b). Filled circle: simulations with the highest similarity to aggregation-stage *Dictyoselium*-rank1 (b, green), HL-60 (b, red) and fish keratocyte (v, orange). (b) The 95% confidence ellipses of the trained datasets (aggregation-stage *Dictyostelium* dark green (dark green), HL-60 (dark red) and fish keratocyte (yellow) in Fig. 1b (a, b) are shown for reference. (c-k) Snapshots and the feature scores of the top-ranked simulations; aggregation-stage *Dictyostelium* (c-e), HL-60 (f-h) and fish keratocyte (i-k).

Although similarity was evaluated based on still images, dynamics of high ranking simulations were by and large consistent with those of real cells. Both *Dictyostelium* and HL60 (Fig 4a,b left panels) showed anterior projections (Fig. 4e bottom panel red regions) that bifurcated from time to time and traveled towards the high curvature region at the rear (*42*). The wave-like appearance was somewhat more prominent in *Dictyostelium*. In both cell types, the posterior end was characterized by a high curvature region that persisted over time. These dynamical features were well recapitulated in the simulations (Fig. 4a,b right panels). Moreover, there was a good agreement between the simulations and the real data in the cell trajectories. The mean square displacement (MSD) of the centroid showed a characteristic time-scale dependency where it was proportional to the square of the elapsed time (*ΔT^2^*) for *ΔT* < *τ*_0_ (Fig. 4c, magenta line) and to *ΔT* for *ΔT* > *τ*_0_ (time domain, Fig. 4c red line). In other words, cells moved ballistically i.e. at a constant velocity for *ΔT* < *τ*_0_ and more like a Brownian particle for *ΔT* > *τ*_0_. *τ*_0_ can be interpreted as the persistence time for directional migration, and square root of the MSD at the inflection point *X*_0_ characterizes the persistence length. Throughout this paper, we chose the time-scale factor τ’= 10 based on approximate matching in the crossover point of the two regression lines between the top ranking simulations and the real-cell data (Fig. 4c,d; red lines). For the top Score-D simulations, we obtained *τ*_0_ = 87 sec, *X*_0_ = 9.8 cell length, compared to *τ*_0_ = 151 sec and *X*_0_ = 15.2 cell-length in the real data which are in good agreement with values reported earlier (*43*). Trajectories of top ranking simulations for Score-H were more persistent (*τ*_0_ = 257 sec, *X*_0_ = 22.1 cell length) as was the case for the real HL60 data (*τ*_0_ = 278 sec, *X*_0_ = 47.7 cell length).

**Figure 4.**
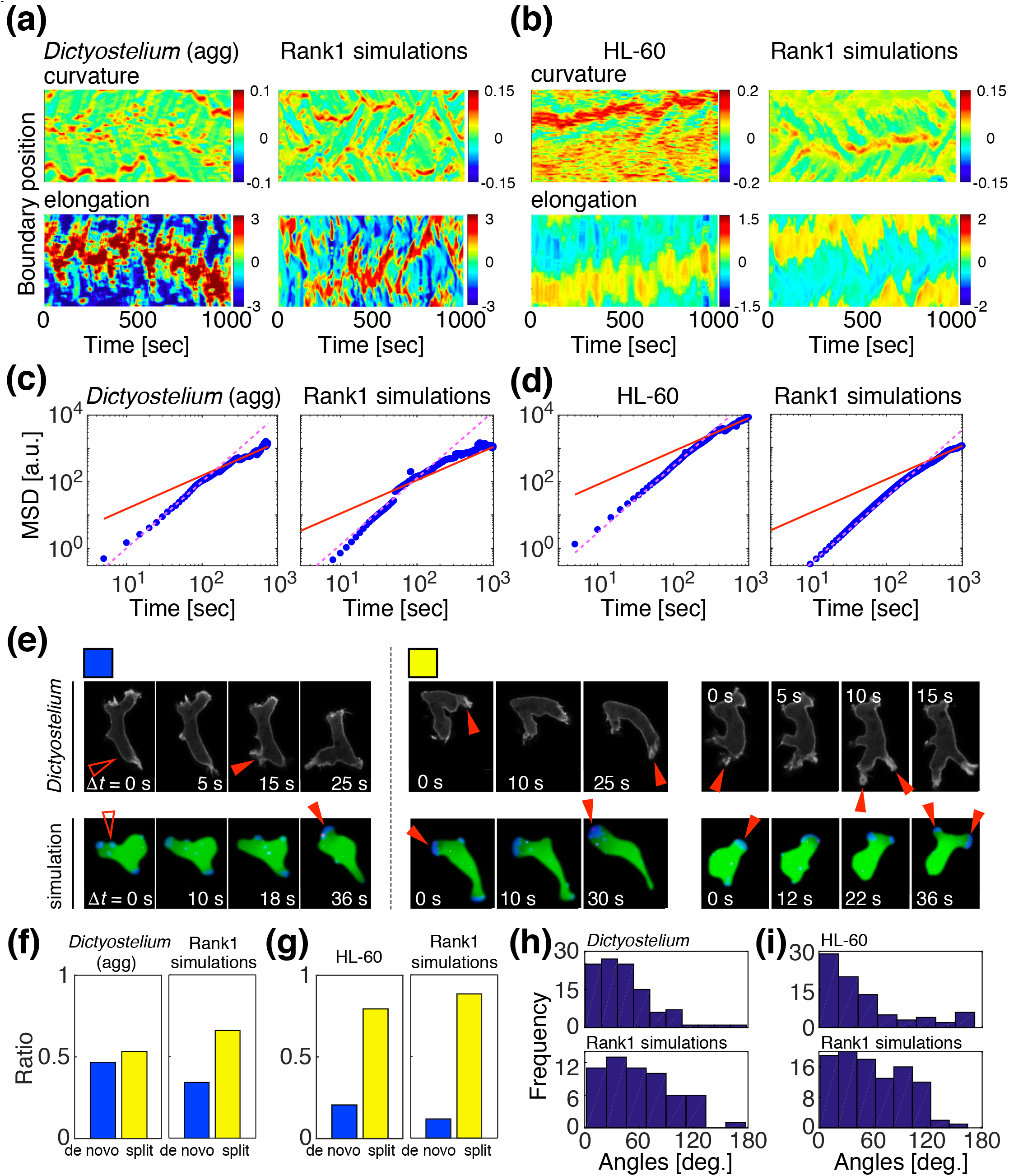
Consistent exploratory dynamics underlie high similarity in the simulated morphologies. (a,b) Local boundary curvature (top panels) and the protrusion speed (bottom panels) (*42*) from the real cell data (left panels) and the top ranked simulations (right panels) for *Dictyostelium* (agg) (a) and HL-60 (b). (c,d) Mean square displacement of the centroid of *Dictyostelium* (agg) (c) and HL-60 (d); real cell data (left panels) and top ranked simulations (right panels). The length scale was normalized by the mean cell length in the moving direction. (e) Representative snapshots of pseudopod formation; de novo formation (left panel) and splitting of an existing pseudopod (one-way-split (middle panel) and Y-shape (right panel)). (f,g) Fractional occurrence of pseudopods by de novo formation (blue) and splitting (yellow); *Dictyostelium* (agg) (f) and HL-60 (g). (h-i) Histogram of pseudopod angles obtained from the time-series of real cell data (a-d left panels) and the top ranked simulations for *Dictyostelium* (agg) (h) and HL-60 (i).

The other important feature of random migration is the relation between pseudopod dynamics and the cell orientation (*7*). New pseudopods frequently appeared in vacant regions (Fig. 4e left), or on top of a pre-existing pseudopod thereby giving the cell the appearance of Y-shape (‘Y-split’ in ref (*7*)). In other cases, pseudopods continued to extend while turning (‘one-way-split’ in ref (*7*)). For aggregation-stage *Dictyostelium* data, the relative occurrence of de novo formation and splitting of pseudopods was approximately 47% and 53% respectively (Fig. 4f). In high ranking simulations, they were 34% and 66%. Similarly for HL60, de novo formation and splitting was 21% and 79% in the top ranking simulation, and in real cell data they were 12% and 88% (Fig. 4g). The extension angles relative to the direction of centroid displacement was about 20-40° for both *Dictyostelium* and HL60 data and their top-ranking simulation counterparts (Fig. 4h,i). Although there was some overrepresentation of extension angles around 90° in the simulation, the angles above 120° were rare in both real data and simulations. All in all, these results demonstrate that the model, albeit its simplification, is able to recapitulate semi-quantitatively both the persistent random walk behavior and the underlying morphology dynamics in *Dictyostelium* and HL-60 cells.

### Mapping of morphological diversification in the feature space predicts key parameters for state transition

Although the datasets analyzed above showed little overlap with one another in the feature space, it should be noted that these coordinates are by no means singular representation of specific cell-types and species from which the data were obtained. As we saw above, there was a large cell-cell variability in the fish keratocyte data that constituted a distinct manifold in the feature space (Fig. 1b; yellow). Likewise, cell-cell variability was evident in the aggregation-stage *Dictyostelium* cells along the PC2 axis (Fig. 1b; green). To see how changes in cell-intrinsic properties alter their positions in the feature space, data from new experimental conditions expected to alter cell polarity were studied (Fig. 5a). Undifferentiated (vegetative) *Dictyostelium* cells took less elongated shape than the aggregation-stage *Dictyostelium* cells under the same substrate and buffer condition. Their aspect ratio on average was smaller than aggregation-stage *Dictyostelium* but larger than that of HL-60 (Fig. 5b). Accordingly, in the PC1-PC2 space, the vegetative *Dictyostelium* was mapped between aggregation-stage *Dictyostelium* and HL-60 (Fig. 5a; magenta reverse triangles). Model simulations that ranked similar to the vegetative *Dictyostelium* data (Fig. 5c; Movie S9) had small *a*_w_ in common (Table S8). Similarly, we analyzed HL-60 cells treated with microtubule destabilizer nocodazole which is known to strengthen neutrophil cell polarity (*44*, *45*). Nocodazole-treated HL-60 cells showed morphology similar to keratocyte with a somewhat smaller aspect ratio (Fig. 5d;) and were mapped between the non-treated HL-60 and the keratocyte datasets (Fig. 5a; blue triangle). The nocodazole-treated HL-60 cells exhibited shape fluctuations making them wobble which was also observed in the simulations (Movie S10). These features were well represented in the respective simulations that ranked high for shape similarity (Fig. 5e). In addition to high *W*_tot_, high ranking simulations had relatively low *a*_w_ or *k*_w1_ (Table S8). This can be interpreted from the fact that low *a*_w_ prevents a cell from breaking apart at high *W*_tot_, while low *k*_w1_ makes the polarized front less pronounced and more sensitive to noise perturbation.

**Figure 5.**
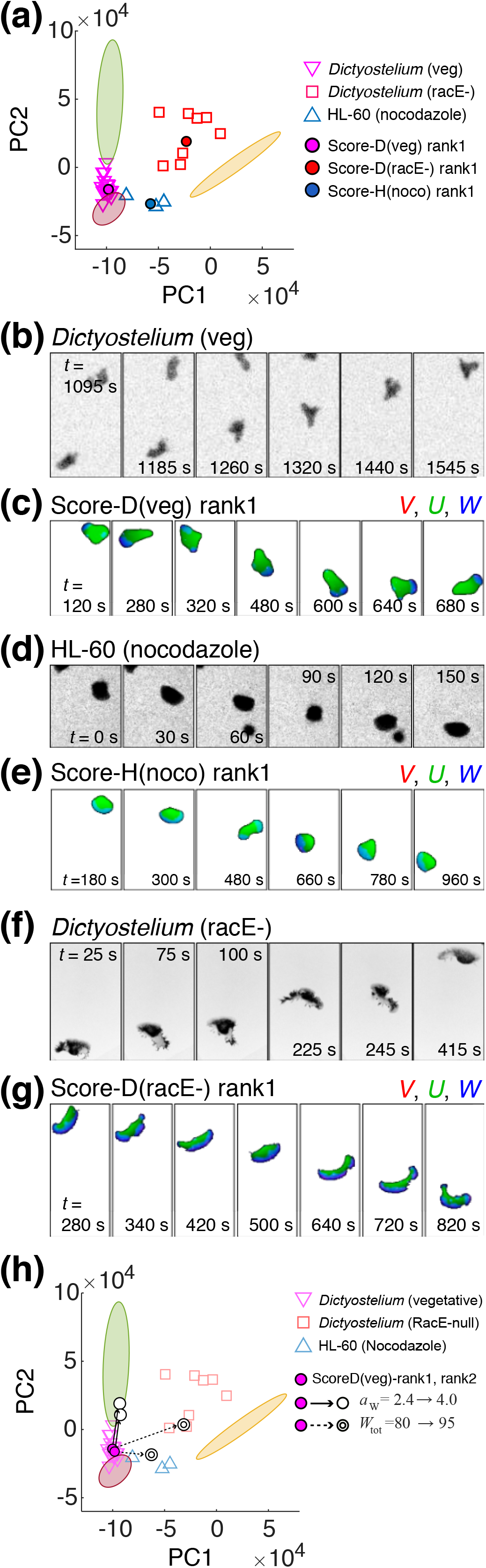
Mapping of non-trained morphologies with perturbing cell polarity predicts a continuous landscape along critical parameters. (a) Morphology features of real cell data from *Dicyostelium* (veg; vegetative stage)(pink inverted triangle), *Dicyostelium* racE- (red square) and nocodazole-treated HL-60 (blue upright triangle) and those from simulations exhibiting highest similarity (filled circles). Colored regions in the background represent the reference microscopy data in Fig. 1b (aggregation-stage *Dictyostelium* dark green (dark green), HL-60 (dark red) and fish keratocyte (yellow)). (b-g) A time-series sample of experimental data closest to the average morphology and the corresponding top-ranked simulations; vegetative-stage *Dictyostelium* (b, c), nocodazole-treated HL-60 (d, e) *Dicyostelium* racE- (f, g). (h) The top-ranked simulations (rank1 and rank2; magenta circles) for *Dicyostelium* (veg) shifted toward *Dictyostelium* (agg) by shifting *a*_W_ from 2.4 to 4 (solid line), and toward the orange region by shifting *W*_tot_ from 80 to 95 (dotted line).

Next, a null strain of *racE (Dicytostelium* RhoA homologue) was chosen for the analysis because of its aberrant cell shape that resembled fish keratocytes (*46*). Under our experimental condition, relatively undifferentiated *racE*- cells exhibited canoe-like shapes that were similar to the fish keratocyte but more dynamic (Fig. 5f). Small fragmented pieces were observed to occasionally split from high curvature regions (Fig. 5f). These data marked relatively high PC1 and PC2 scores and mapped between the keratocyte and the aggregation-stage *Dictyostelium* data (Fig. 5a; black square). The top ranking simulations for the *racE*- data (Table S8; high Score-D (*racE*-)) exhibited remarkably similar morphology dynamics characterized by high lateral deformation and occasional fragmentation (Fig. 5g; Movie S11). As expected, there was a large overlap with high Score-K data, leaving only the top 3 Score-K data (Table S7 bottom rows; ||*F*||_2_ <3 x 10^4^) that uniquely mapped to the keratocyte data. Compared to either the high Score-D (veg) or the high Score-D (agg) data, the high Score-D (*racE-*) data had large *W*_tot_ (Table S8) consistent with the laterally extended cell shape. All in all, the above analysis suggests key parameters *a*_w_ and *W*_tot_ that are pivotal for the state transition between the characteristic morphologies. This was further verified by studying unexamined regions in the high-dimensional parameter space near the top ranking simulations for vegetative-stage *Dictyostelium*. Increasing the value of *a*_w_ brought the morphology score closer to that of the aggregation-stage cell (Fig. 5h). In contrast, increasing *W*_tot_ increased PC1 and brought the shape state closer to the nocodazole-treated HL-60 and *racE*- cells (Fig. 5h).

## Discussion

In this study, we presented a hybrid cell morphology analysis that combined deep-learning-based extraction of morphological metrics from microscopy data and dynamical model simulations. Convolutional neural network was trained to classify stereotype morphology of migrating cells represented by HL-60, *Dictyostelium* and fish keratocytes. The feature vector of the trained classifier showed that the three representative morphologies can be described in low dimensional feature space; the first component represents elongation in the lateral direction and the second component represents anterior-posterior elongation and edges. The new feature space essentially expanded the principle shape variations associated with cell polarity previously identified in the keratocyte data (*18*) to encompass narrowly polarized cell morphologies with pseudopodal extensions. The finding of low dimensional feature space is in line with an earlier feature representation of *Dictyostelium* cells based on Fourier-mode decomposition of the cell contour (*20*). While being less analytically clear than the Fourier analysis, the present approach allows one to distinguish morphology with respect to the cell orientation and thus suited to analyze migratory cells. Interestingly, sample conditions expected to alter cell polarity – vegetative stage *Dictyostelium*, *racE*-/*Dictyostelium*, and the nocodazole treated HL-60 cells were mapped to intermediate coordinates spanned by these two major manifolds. Moreover, the new sample conditions were each found clustered without significant overlaps with the training data sets. These results suggest that the region spanned by the three training datasets (keratocyte, H60, *Dictyostelium* (agg)) in the PC1-PC2 coordinate constitutes a space continuously occupied by forms realizable by genetic and phenotypic variations.

Our signaling network model was able to identify regions in the feature space realizable by fast migrating cells that have not been recapitulated in earlier modeling studies. These include narrowly polarized cell morphology, pseudopods as well as their aberrant forms. Parameters chosen based on the similarity with still images provided morphodynamic basis for the random walk trajectory previously approximated by a particle model (*1*). The main difference between existing models and the present model is that while previous models were either rule-based dynamics (*37*, *47*), excitable and oscillatory dynamics (*14*, *40*, *48*) or the wave-pinning dynamics (*38*, *39*), our work addressed morphological outcomes of the combination of excitable protrusion and bistable polarization that compete for dominance under limited *W*_tot_ i.e. the maximal protrusive activity in addition to a positive feedback regulation between the two. In the model, this is described by the *W*-dependency for the amplification of *V* (Eq. 2c) and amplification of *W* by *V* (Eq. 2b). Although how exactly the coupling is implemented biochemically requires further investigation, in neutrophils, Cdc42 acts globally to enforce cell polarity by promoting actomyosin contractility through its effector WASP in a microtubule dependent manner (*49*). Recent studies indicate that, in *Dictyostelium*, Ras at the leading edge interacts with GDP-bound form of RacE to strengthen cell polarity (*36*). Mapping of the *racE*- morphology to a high *W*_tot_ state (Fig. 5h) hints at the nature of the competition for *W*_tot_ in respect to the states of actin: the contractile cortical meshwork that are crosslinked with myosin II and the protrusive dendritic meshwork that requires the Arp2/3 complex for side-branching nucleation. RacE is essential for plasma membrane localization of Diaphanous-related formins (DRFs) (*46*), and deletion of DRFs (ForA-/ForE-/ForH-) results in the loss of cortical actin. Since the morphology of the null mutant of DRFs phenocopies that of the *racE*- null cells (*46*), the increase in *W*_tot_ are associated with the absence of DRFs from the plasma membrane. On the other hand, the fluctuating protrusions are largely associated with fast idling pulses of Scar/Wave activities which are amplified by the excitable network (*50*). Recent studies suggested that actin nucleators such as formins and Arp2/3 are competing for a limited pool of actin monomers and/or their upstream activators such as Rac-GTP (*51–53*). Such notion is also supported by an observation that the amount of F-actin is compensated in Scar-/WASP-cells by increased localization of ForH at the cortex (*54*). Taken together with our mapping of *racE-* data, these observations are in line with our current model view that excitability and cell polarity networks compete for dominance over limited *W*_tot_.

The variations in the distinct morphologies of differentiating *Dictyostelium* cells suggest alterations in the key parameters that serves as a control point. The difference between the vegetative- and aggregation-stage *Dictyostelium* was ascribed mainly to an increase in the membrane protrusion force *a*_w_. The increase in *a*_w_ can be understood from the fact that Rac1 (*55*) and SCAR (*56*), the essential factors for Arp2/3 activation, are known to be expressed at low levels in the vegetative-stage then increase markedly in the aggregation-stage cells. A recent study based on an excitable model (*40*) suggested a progressive state transition from a circular to amoeboid then to a keratocyte-like shape by the increase in the protrusion force (Fig. 2f. in (*40*)). Rather, our model predicts that changes in the protrusive force should allow a direct transition from a circular shape (low PC1 low PC2) state to either amoeboid or keratocyte-like form depending on *W*_tot_. Such direct transition has been demonstrated experimentally and was attributed to an increased activity of a nested excitable network (*14*). However, the elongated shapes in their model were oscillatory and lacked the persistency. The presence of the polarity dynamics underlies the characteristic longitudinally extended cell shapes accompanied by branching pseudopod which do not arise in these models(*14*, *40*). Relatively low *D*_w_ is required in both vegetative and aggregation-stage *Dictyostelium* to prevent the polarity dynamics from completely winning over the excitable dynamics. A highly polarized form at high *D*_w_; i.e. the 1-variable model limit *ζ*/ *D*_w_ → 0 (fig. S3e-h; Supplementary Text) is indeed reached by cells that further differentiated into prespore cell-type (fig. S3i). While it is possible that certain cells are in a decoupled state (*ζ* = 0), the requirement of large *D*_w_ for cell polarity signifies the importance of *W* acting globally. Large *D*_w_ may not necessarily be mediated by pure diffusion as the present model postulates, but instead could be realized by other transport processes implicated in cell polarity such as membrane flow or the myosin-II dependent global actin flow. Global actin flow has been shown to maintain asymmetric distribution of de-filamenting factors (*57*, *58*), however such global flow may not always be present in polarized cells (*59*). Since these transport processes are tied to cortical actin, they are naturally accompanied by changes in membrane tension (*60*) which should also be part of the feedback process from *W* to *V*.

The present framework of data analysis potentially provides means to test and improve specific models of migrating cells. Distinctions between various excitable (*14*, *41*, *61*) and cell polarity models (*38*, *62*, *63*) will become more relevant as we proceed further to analyze detailed geometries and dynamics associated with specific cells and conditions. For example, our ideal cell model gave more frequent rise to pseudopods from the tail region compared to the real cell data. Such a discrepancy could be due to the fact that retraction is assumed to be driven only by the area conservation and that no regulated contractility was explicitly described. While this approximation can be justified when there is reciprocity between the front expansion and the rear contraction as has been shown to hold independently of actomyosin in neutrophils (*45*), the present model could be modified in the future to include local cortical actomyosin regulation when analyzing detailed shapes of the cell rear and the bleb-based front protrusion. Further improvement of the model and increasing dimensionality of the feature space may work hand in hand with extending the present analysis to classify morphologies exhibited by other cell types of wildtype and mutant backgrounds. For example, the present analysis fails to distinguish the pancake-like shape known for Rac and Rap related mutants that result from uniform expansion (*41*) and similarly round (i.e. low PC1, low PC2) cells inhibited of actin polymerization. This limit maybe overcome by introducing absolute size instead of normalizing the area so as to distinguish spherical cell versus flattened cell in two-dimensional cell masks. Expanding the analysis to 3-dimensional images would also be better suited to the present machine learning approach. As resolutions and dimensions are increased, the cell-shape based analysis may be supplemented with fluorescence image data of cytoskeletons and their regulators. Given the significant bottleneck in the present simulation by the huge computational loads which required parallel computation by GP-GPU, other avenues of coarse-graining maybe required to extend the present approach to a larger multi-modal analysis.

## Materials and Methods

### Dictyostelium and HL-60 cell culture and data acquisition

Time-lapse data of freely migrating *Dictyostelium*, neutrophil-like HL-60, fish keratocyte and differentiated T mouse cells were acquired with an inverted microscope using either 20, 40 or 60x objective lens. For *Dictyostelium* and HL-60, cells expressing Lifeact fused to mNeonGreen (*64*) and mTurquoise2 were employed, respectively. A Lifeact-mTurquoise2 expression vector was constructed by ligating Lifeact-mTurquoise2 into an episomally replicating plasmid pEBMulti Neo (WAKO, 057-08131) at restriction sites *Xho*I and *Not*I. The Lifeact-mTurquoise2 expressing stable HL60 cell line was obtained by introducing the plasmid by electroporation (NEPA21; Nepa Gene, Ltd., Chiba, Japan) followed by G418 selection (1 mg mL^−1^) after 2 days. For fish keratocytes, DIC images were employed. *Dictyostelium* cells were grown axenically and obtained according to standard protocols as previously described (*65*). Vegetative *Dictyostelium* AX4 cells (N = 1694 snapshots from 18 timeseries;), aggregation-stage *Dictyostelium* AX4 cells (starved for 3.5 hours; N = 2841 snapshots 19 timeseries;), vegetative LifeactGFP/*rac*E- cells (N = 330 snapshots from 8 timeseries;) were plated on a non-coated coverglass and images were acquired at 5 sec/frame (aggregation-stage AX4 and *rac*E- cell) or 15 sec/frame (vegetative AX4). Neutrophil-like HL-60 cells were grown in RPMI1640/glutamate media (Wako 189-02145) supplemented with 12% FBS (Sigma 172012). Differentiated HL-60 cells were obtained by treating the cells with DMSO for 3 days. Images of HL-60 cells on fibronectin-coated glass plates in the presence of 1 nM fMLP (N = 3468 snapshots from 23 timeseries;) were taken at 5 sec/frame. For nocodazole treatment, differentiated HL-60 cells were collected by centrifugation, suspended in fresh HBSS containing 20 μM nocodazole and plated on a coverslip pre-coated with 1-2% BSA in PBS. Data were acquired within 20-75 min in the same medium in the presence of nocodazole (N = 181 snapshots from 3 timeseries).

### Primary cell culture and data acquisition

Keratocytes from the scales of Central American cichlids (*Hypsophrys nicaraguensis*) were cultured as previously described (*66*) and images were recorded at 2-s intervals (N = 1590 snapshots from 12 timeseries;). Naïve CD4^+^ T cells were isolated from the lymph nodes and spleen of C57BL/6 mice by MidiMACS (Miltenyi Biotec). Cells were activated by plate-bound anti-CD3 (5 μg mL^−1^) and anti-CD28 (2.5 μg mL^−1^) with cytokines and blocking antibodies. Th1: 2 ng mL^−1^ hIL-2, 5 ng mL^−1^ mIL-12. Treg: 1 ng mL^-1^ hTGFβ, 1 μg mL^−1^ anti-IFNγ. Snapshots from these timelapse recording were employed for feature extraction.

### Deep-learning-based feature extraction

Mask images were pre-processed as follows (Fig. 1a, left panel): (i) the migration direction was determined from the centroid displacement at a five timeframe interval (equivalent to 25 sec for aggregation-stage and *rac*E- *Dictyostelium* data, 10 sec for keratocyte data) except for vegetative *Dictyostelium* data where 1 timeframe (equivalent to 15 sec) was used, (ii) binarized mask image was rotated to align the migration direction to the y-axis, (iii) the image was rescaled so that the cell area is equal to the area of a circle with 25 pixel diameter. The rescaled masks were each embedded at the center of a blank square frame of 64×64 pixels. The exact spatial resolution of mask images varied from sample to sample due to rescaling, however they were all in the order of ~0.5 μm/pixel. Convolutional neural network (Figure 1a, bottom panel) was implemented using Keras (https://keras.io) with TensorFlow backend. To make the sample size of the three datasets near equal, data augmentation was performed by rotating the original masks at angles (< ±5 deg) randomly picked from a uniform distribution (see Table 1 for the number of samples). Input vectors were processed through layers of convolution operation (Fig. 1a, bottom panel; ‘convolution’) in addition to layers of max pooling operation with a 3×3 kernel to render the analysis robust to positional deviation (Fig. 1a, bottom panel; ‘pooling’). These were then processed through a set of densely connected layers with rectified linear and hyperbolic tangent activation function (Fig. 1a, bottom panel; ‘rel’ and ‘tanh’). In the final layers, the dimension of the vector was reduced to three and were passed to ‘softmax’ activation function. The values of the three nodes (*F*_1_, *F*_2_, *F*_3_) before the final softmax layer were employed to represent cell shapes. The number of training epoch was 2000 which was sufficient for adequate learning as determined by the accuracy and loss values (fig. S1a,b).

### Geometrical analysis of feature space

To examine mapping of 2-dimensional geometries in the feature space, 7 well-defined objects were used; circle, isosceles triangle, asymmetric right triangle, rectangle, rhombus, pentagon and star polygon in a specified orientation (Fig. 1n). Triangles, pentagon and star were flipped vertically to obtain total of 11 basic objects. The library was further expanded to 33 shapes by including variants of the basic geometries with different aspect ratio 1:1, 1:2 (Fig. 1n red, horizontally elongated shape to the moving direction) and 2:1 (Fig. 1n blue, flattened shape with vertically elongated to the moving direction).

### Model Equations

To numerically simulate the interface between the plasma membrane and the extracellular space, we employed a phase-field equation in the following form (*38*, *62*, *67*).

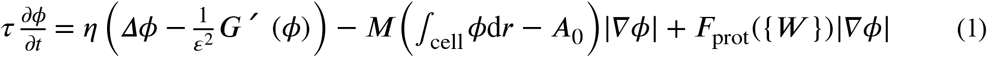

The equation describes the dynamics of a continuous state variable *ϕ*(***r***; *t*) in a two-dimensional space that specifies whether a position ***r*** is occupied (*ϕ* = 1) or not occupied (*ϕ* = 0) by a cell at time *t*. In the present study, we consider initial conditions with a single continuous domain with *ϕ* = 1. The parameter *τ* is the viscous friction coefficient. The first term of the r.h.s represents effective surface tension, where Δ*ϕ* and *G* are derived from membrane energy. ***G*** is Landau functional describing a bi-stable potential. Here we chose *G*(*ϕ*) = 18*ϕ*^2^(1 – *ϕ*)^2^ (62). The second term describes restoring force that keeps the cell area close to *F*_prot_ The Eq. 1 assumes that the bending energy is negligible (*67*). The third term represents active force with magnitude *F*_prot_ that is perpendicular to the boundary |∇*ϕ*| ≠ 0 and thus drives membrane extension. In the present simulations, parameters in Eq. 1 were set so that they are an order of magnitude within generally accepted values (Table S2); surface tension *η* = 1.0 [pN], (*68*), cell area *A*_0_ = 78.83 [μm^2^] (~ 5 μm radius circle), protrusive force by actin polymerization *a*_w_*W* = 0.8 – 4.0 [pN/μm] (for *W* ~ 1) (*69*, *70*). Since *τ* was not well constrained experimentally and expected to differ between cell-types and the culture conditions, we adopted an empirical value 0.83 [pN/μm^2^] (*67*), which was then calibrated retrospectively by a multiplier *τ’* so that the time scale of the simulated cell trajectories match with that of real data as described in the later section. The area constraint *M* = 0.5 pN/μm^3^ and the size of the boundary layer *ε* = 1.0 [μm] were set close to those in the earlier studies (*62*, *67*).

The ‘ideal cell’ that can take close to all of the basic phases of morphology features that we have examined (Fig. 1) should consist of two main features: 1) transient appearance of localized protrusions and 2) prolonged presence of single expanding edge and retracting tail to appear under homogeneous extracellular conditions (*1*). Here, we formulate a reaction-diffusion model that describes these two processes mathematically as follows:

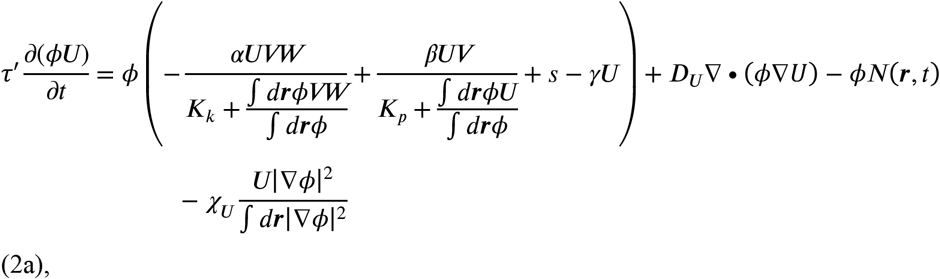

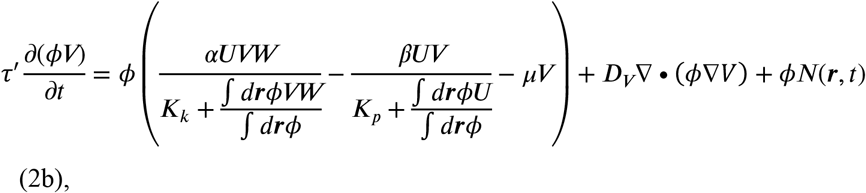

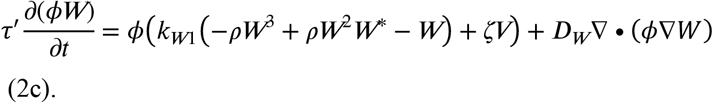

where the first two equations for *U* and *V* describe an excitable reaction network for transient protrusive dynamics, and the third equation for *W* describes polarization dynamics, respectively (Fig. 2a; Supplementary Text). The role of the excitable reaction network is to generate transient signals for local protrusions in a minute to a few minutes time-scale by amplifying small edge fluctuations of seconds order (*50*). In neutrophils, excitable dynamics of Cdc42 and PI3K activity (*32*, *33*) is essential for front protrusions. In *Dictostelium*, excitable dynamics are observed at the level of spontaneous Ras and PI3K activation (*30*, *67*, *71*). Here, we adopted equations originally introduced to study excitable PI3K activities and the resulting F-actin waves in *Dictyostelium* (Eq. 2a and 2b) (*67*). Parameters *α* and *β* are the rate constants of reaction *U* → *V* and *V* → *U* multiplied by the time-scale factor *τ’*. The source of edge fluctuation is introduced as a noise term *ϕN*(***r***,*t*) (*67*) in Eq. 2b which is amplified through *V* by a positive-feedback described in the first term. Increase in *V* is then slowed down due to depletion in *U*, and the system eventually recovers the original resting state.

The expanding membrane region is determined by *W* governed by Equation (2c) (Fig. 2a; see Supplementary Text for derivation) which is similar in form to the wave-pinning (*29*). The same type of equation has been used earlier to study polarized cell shape in fish keratocyte (*63*) and *Dictyostelium* (*38*). The first term describes reaction kinetics with bistability, and the second term describes diffusion. We assume that the sum of *W* and its reciprocal state *W*\* is conserved (*W*_tot_ = const.) (*29*) and that *W*\* diffuses much faster than *W* and thus can be approximated as uniform in space. Hence we obtain_

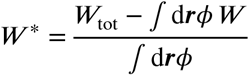

Due to the global constraint, this class of bistable system gives rise to coexistence of low and high *W* regions due to stalling of a transitory wave front - a property known as “wave pinning” (*29*, *63*). For the sake of brevity, we shall embed rear retraction passively in the form of restoring force. Thus we set *F*_prot_ = *a*_W_*W* in Eq. 1 (*62*) so that the edge expands where *W* is high, and the rest of the domain with high *W*_tot_ /*A*_0_ -*W* as a result contracts due to area conservation imposed by the second term in Eq. 1. Note that *W* only specifies where the protrusions and retraction take place and does not assume the origin of their driving force. The variable *W* can thus be interpreted as an abstract representation of bistable signals such associated with the leading edge of polarize cells as Rac-GTP, or alternatively, *W*_tot_/*A*_0_ - *W* with the reciprocal spatial profile to indirectly represent signals that regulate rear contraction such as RhoA-GTP. A recent study has shown that Rac-GTP accumulates at the front of migrating neutrophils regardless of whether it is F-actin filled or blebbing (*60*). Based on the inherent ability to maintain persistent front-and-back unipolarity, the polarity network (Eq. 2c) serves as a spatio-temporal filter to select or remove local protrusions. This coupling is described by assuming that *W* positively regulates the positive-feedback amplification of *V* (Eq. 2c; r.h.s first term) in addition to *W* feeding back to increase *V* (Eq. 2a) to reflect F-actin- or tension-dependent positive regulation of leading edge signals in neutrophils and *Dictyostelium* (*60*, *72*, *73*).

### Parameter search

Model parameters were selected for the systematic feature analysis (Fig. 3a; Table S6) based on the following criteria: Of the total 22 parameters (Table S2 and S3), eight parameters (*W*_tot_, *a*_W_, *k*_W1_, *μ, g, ρ, D*_w_, *χ*_U_) were chosen for detail analysis. Parameters (*a, β, K_k_, K_P_, s*) that define the kinetics of *U*, *V* were fixed to those used earlier (*67*). Two to four independent simulations from different random seeds were executed for a given parameter set to obtain average feature scores ***F***. Because grid-search in 8 dimensional parameter space was unattainable due to heavy computational load, we adopted parameter search around manually selected reference parameter sets. First, based on preliminary simulations performed with various parameters, a single parameter set R1 was chosen which gave rise to polarized cell morphology; i.e. elongated shape in the moving direction. Around the reference set R1, the following parameters were varied: *a*_w_ and *W*_tot_ which appeared to affect the elongation of cell in the moving direction, *X* and *D*_W_ related to the split of the leading edge, and *γ* that seems to affect the appearance of new pseudopods. We performed a grid search around R1 in 2-dimension (*a*_W_, *g*) for *χ*_U_ = 0 and *χ*_U_ =50 (fig. S4a left panels) while fixing other 6 parameters. Similarly, a grid search was performed in (*W*_tot_, *D*_W_) for *χ*=0 and 50 (fig. S4a left panels). Second, we chose another parameter set R2 which gave rise to relatively round shape and random cell displacement (i.e. near HL60-like behavior). Around R2, we varied (*a*_W_, *W*_tot_), (*g*, *ρ*), and (*k*_W1_, *μ*) at both *χ*_U_ = 0 and 50 (fig. S4a right panels). Representative results of the simulations are shown in figure S4b and c. Phase diagram of feature vector ***F*** around R1 on the *γ*- *a*_W_ plane and around R2 on *μ* - *k*_W1_ plane are shown in figure S4d and e, respectively.

### Pseudopod analysis

For both experiments and simulation data, events of pseudopod formation were detected manually by eye as either ‘de novo’, ‘Y-split’ or ‘one-way-split’ based on ref. (*7*). New pseudopods that appeared in non-protruding regions (Fig. 4e left) were counted as “de novo formation”. Protrusions that appeared close to the leading edge were marked as ‘Y-split’ because of the resulting cell shape (Fig. 4e right). When new protrusive events occurred on top of the leading edge so as to offset and steer the extending pseudopod, it was counted as ‘one-way-split’ (Fig. 4e middle) (*7*). For the subset of the data, the analysis was performed three independent times to confirm reproducibility. To obtain angular distribution, direction of pseudopod extension 10 seconds from the onset was measured relative to the centroid movement. Aggregation-stage *Dicyostelium* (N = 90 from 4 timeseries), and the closest simulations (N = 44 from 2 timeseries). HL-60 (N = 68 from 5 timeseries), and the closest simulations (N = 77 from 3 timeseries).

## Acknowledgements

The authors thank Shuji Ishihara for discussion and helpful comments. HL-60 cells were obtained from BRC Cell Bank (RCB0041, RIKEN). Lifeact-mTurquoise2 was a gift from Dorus Gadella (Addgene plasmid #32601). The authors are also grateful to Hiroshi Senoo and Miho Iijima (Johns Hopkins University) for the *rac*E- cells, Jonathan Chubb (University College London) for the *Dictyostelium* codon optimized NeonGreen, Douwe M. Veltman (University of Groningen) for the *Dictyostelium* expression vector pDM. This work was funded by grants from Japan Science and Technology Agency (JST) CREST JPMJCR1923, Japan Society for Promotion of Science (JSPS), Ministry of Education, Culture, Sports, Science and Technology (MEXT) KAKENHI JP19H05801 to SS and in part by Joint Research by Exploratory Research Center on Life and Living Systems (ExCELLS) Grant 18-204, MEXT KAKENHI JP19H05416, JP18H04759 and JP16H01442; JSPS KAKENHI JP17H01812 and JP15KT0076 (to S.S.).

## References

1. L. Li, S. F. Nørrelykke, E. C. Cox, Persistent cell motion in the absence of external signals: a search strategy for eukaryotic cells. PLoS ONE. 3, e2093 (2008).

2. M. Otsuji, Y. Terashima, S. Ishihara, S. Kuroda, K. Matsushima, A conceptual molecular network for chemotactic behaviors characterized by feedback of molecules cycling between the membrane and the cytosol. Sci Signal. 3, ra89–ra89 (2010).

3. X. Liu, E. S. Welf, J. M. Haugh, Linking morphodynamics and directional persistence of T lymphocyte migration. J R Soc Interface. 12, 20141412–20141412 (2015).

4. M. F. Krummel, F. Bartumeus, A. Gérard, T cell migration, search strategies and mechanisms. Nat Rev Immunol. 16, 193–201 (2016).

5. C. M. Witt, S. Raychaudhuri, B. Schaefer, A. K. Chakraborty, E. A. Robey, Directed Migration of Positively Selected Thymocytes Visualized in Real Time. PLoS Biol. 3, e160–8 (2005).

6. N. Andrew, R. H. Insall, Chemotaxis in shallow gradients is mediated independently of PtdIns 3-kinase by biased choices between random protrusions. Nat Cell Biol. 9, 193–200 (2007).

7. L. Bosgraaf, P. J. M. Van Haastert, The ordered extension of pseudopodia by amoeboid cells in the absence of external cues. PLoS ONE. 4, e5253 (2009).

8. A. Leithner et al., Diversified actin protrusions promote environmental exploration but are dispensable for locomotion of leukocytes. Nat Cell Biol. 18, 1253–1259 (2016).

9. L. K. Fritz-Laylin et al., Actin-based protrusions of migrating neutrophils are intrinsically lamellar and facilitate direction changes. Elife. 6, 437 (2017).

10. M. Bergert, S. D. Chandradoss, R. A. Desai, E. Paluch, Cell mechanics control rapid transitions between blebs and lamellipodia during migration. Proc Natl Acad Sci USA. 109, 14434–14439 (2012).

11. R. J. Petrie, A. D. Doyle, K. M. Yamada, Random versus directionally persistent cell migration. Nat Rev Mol Cell Biol. 10, 538–549 (2009).

12. J. Lee, Insights into cell motility provided by the iterative use of mathematical modeling and experimentation. AIMS Biophysics. 5, 97–124 (2018).

13. Y. Asano et al., Keratocyte-like locomotion in amiB-null Dictyostelium cells. 59, 17–27 (2004).

14. Y. Miao et al., Altering the threshold of an excitable signal transduction network changes cell migratory modes. Nat Cell Biol. 19, 329–340 (2017).

15. Z. Chen et al., Gleevec, an Abl Family Inhibitor, Produces a Profound Change in Cell Shape and Migration. PLoS ONE. 8, e52233–14 (2013).

16. A. Van Keymeulen et al., To stabilize neutrophil polarity, PIP3 and Cdc42 augment RhoA activity at the back as well as signals at the front. J Cell Biol. 174, 437–445 (2006).

17. H. Hattori et al., Small-molecule screen identifies reactive oxygen species as key regulators of neutrophil chemotaxis. Proc Natl Acad Sci USA. 107, 3546–3551 (2010).

18. K. Keren et al., Mechanism of shape determination in motile cells. Nat. Cell. Biol. 453, 475–480 (2008).

19. X. Ma, O. Dagliyan, K. M. Hahn, G. Danuser, Profiling cellular morphodynamics by spatiotemporal spectrum decomposition. PLoS Comput Biol. 14, e1006321 (2018).

20. L. Tweedy, B. Meier, J. Stephan, D. Heinrich, R. G. Endres, Distinct cell shapes determine accurate chemotaxis. Sci. Rep. 3, 2606 (2013).

21. E. Alizadeh, S. M. Lyons, J. M. Castle, A. Prasad, Measuring systematic changes in invasive cancer cell shape using Zernike moments. Integr Biol (Camb). 8, 1183–1193 (2016).

22. M. K. Driscoll, J. T. Fourkas, W. Losert, Local and global measures of shape dynamics. Phys Biol. 8, 055001 (2011).

23. R. M. Cooper, N. S. Wingreen, E. C. Cox, An excitable cortex and memory model successfully predicts new pseudopod dynamics. PLoS ONE. 7, e33528 (2012).

24. C. Bakal, J. Aach, G. Church, N. Perrimon, Quantitative morphological signatures define local signaling networks regulating cell morphology. Science. 316, 1753–1756 (2007).

25. Z. Yin et al., A screen for morphological complexity identifies regulators of switch-like transitions between discrete cell shapes. Nat. Cell. Biol. 15, 860–871 (2013).

26. G. Danuser, J. Allard, A. Mogilner, Mathematical modeling of eukaryotic cell migration: insights beyond experiments. Annu Rev Cell Dev Biol. 29, 501–528 (2013).

27. V. Te Boekhorst, L. Preziosi, P. Friedl, Plasticity of Cell Migration In Vivo and In Silico. Annu Rev Cell Dev Biol. 32, 491–526 (2016).

28. J. Lee, A. Ishihara, J. A. Theriot, K. Jacobson, Principles of locomotion for simple-shaped cells. Nat. Cell. Biol. 362, 167–171 (1993).

29. Y. Mori, A. Jilkine, L. Edelstein-Keshet, Wave-pinning and cell polarity from a bistable reaction-diffusion system. Biophys J. 94, 3684–3697 (2008).

30. P. J. M. Van Haastert, I. Keizer-Gunnink, A. Kortholt, Coupled excitable Ras and F-actin activation mediates spontaneous pseudopod formation and directed cell movement. Mol Biol Cell. 28, 922–934 (2017).

31. C. Arrieumerlou, T. Meyer, A Local Coupling Model and Compass Parameter for Eukaryotic Chemotaxis. Dev Cell. 8, 215–227 (2005).

32. H. W. Yang, S. R. Collins, T. Meyer, Locally excitable Cdc42 signals steer cells during chemotaxis. Nat Cell Biol. 18, 191–201 (2015).

33. M. Tang et al., Evolutionarily conserved coupling of adaptive and excitable networks mediates eukaryotic chemotaxis. Nat Commun. 5, 5175 (2014).

34. K. Plak et al., GxcC connects Rap and Rac signaling during Dictyostelium development. BMC Cell Biol. 14, 6 (2013).

35. S. Beco et al., Optogenetic dissection of Rac1 and Cdc42 gradient shaping. Nat Commun, 1–13 (2018).

36. H. Senoo et al., Phosphorylated Rho-GDP directly activates mTORC2 kinase towards AKT through dimerization with Ras-GTP to regulate cell migration. Nat Cell Biol. 21, 867–878 (2019).

37. I. Niculescu, J. Textor, R. J. de Boer, Crawling and Gliding: A Computational Model for Shape-Driven Cell Migration. PLoS Comput Biol. 11, e1004280 (2015).

38. S. Alonso, M. Stange, C. Beta, Modeling random crawling, membrane deformation and intracellular polarity of motile amoeboid cells. PLoS ONE. 13, e0201977 (2018).

39. W. R. Holmes, A. E. Carlsson, L. Edelstein-Keshet, Regimes of wave type patterning driven by refractory actin feedback: transition from static polarization to dynamic wave behaviour. Phys Biol. 9, 046005 (2012).

40. Y. Cao, E. Ghabache, W.-J. Rappel, Plasticity of cell migration resulting from mechanochemical coupling. Elife. 8, 107 (2019).

41. M. Edwards et al., Insight from the maximal activation of the signal transduction excitable network in Dictyostelium discoideum. Proc Natl Acad Sci USA. 115, E3722–E3730 (2018).

42. M. K. Driscoll et al., Cell shape dynamics: from waves to migration. PLoS Comput Biol. 8, e1002392 (2012).

43. H. Takagi, M. J. Sato, T. Yanagida, M. Ueda, PLOS ONE: Functional Analysis of Spontaneous Cell Movement under Different Physiological Conditions. PLoS ONE (2008).

44. J. Xu, F. Wang, A. Van Keymeulen, M. Rentel, Neutrophil microtubules suppress polarity and enhance directional migration. Proc Natl Acad Sci USA. 102, 6884–6889 (2005).

45. T. Y.-C. Tsai et al., Efficient Front-Rear Coupling in Neutrophil Chemotaxis by Dynamic Myosin II Localization. Dev Cell. 49, 189–205.e6 (2019).

46. C. Litschko et al., Functional integrity of the contractile actin cortex is safeguarded by multiple Diaphanous-related formins. Proc Natl Acad Sci USA. 116, 3594–3603 (2019).

47. J. Satulovsky, R. Lui, Y.-L. Wang, Exploring the control circuit of cell migration by mathematical modeling. Biophys J. 94, 3671–3683 (2008).

48. O. Nagel et al., Geometry-Driven Polarity in Motile Amoeboid Cells. PLoS ONE. 9, e113382 (2014).

49. S. Kumar et al., Cdc42 regulates neutrophil migration via crosstalk between WASp, CD11b, and microtubules. Blood. 120, 3563–3574 (2012).

50. C.-H. Huang, M. Tang, C. Shi, P. A. Iglesias, P. N. Devreotes, An excitable signal integrator couples to an idling cytoskeletal oscillator to drive cell migration. Nat Cell Biol. 15, 1307–1316 (2013).

51. T. A. Burke et al., Homeostatic Actin Cytoskeleton Networks Are Regulated by Assembly Factor Competition for Monomers. Current Biology. 24, 579–585 (2014).

52. A. J. Lomakin et al., Competition for actin between two distinct F-actin networks defines a bistable switch for cell polarization. Nat. Cell. Biol. 17, 1435–1445 (2015).

53. C. Suarez, D. R. Kovar, Internetwork competition for monomers governs actin cytoskeleton organization. Nat Rev Mol Cell Biol. 17, 799–810 (2016).

54. A. J. Davidson, C. Amato, P. A. Thomason, R. H. Insall, WASP family proteins and formins compete in pseudopod-and bleb-based migration. J Cell Biol. 217, 701–714 (2018).

55. J. Bush, K. Franek, J. Cardelli, Cloning and characterization of seven novel Dictyostelium discoideum rac-related genes belonging to the rho family of GTPases. Gene. 136, 61–68 (1993).

56. J. E. Bear, J. F. Rawls, C. L. Saxe, SCAR, a WASP-related Protein, Isolated as a Suppressor of Receptor Defects in Late Dictyostelium Development. J Cell Biol. 142, 1325–1335 (1998).

57. P. Maiuri et al., Actin Flows Mediate a Universal Coupling between Cell Speed and Cell Persistence. Cell. 161, 374–386 (2015).

58. L. Yolland et al., Persistent and polarized global actin flow is essential for directionality during cell migration. Nat. Cell. Biol., 1–27 (2019).

59. Y. Fukui, T. Kitanishi-Yumura, S. Yumura, Myosin II-independent F-actin flow contributes to cell locomotion in dictyostelium. J Cell Sci. 112 (Pt 6), 877–886 (1999).

60. B. R. Graziano et al., Cell confinement reveals a branched-actin independent circuit for neutrophil polarity. PLoS Biol. 17, e3000457–34 (2019).

61. F. Knoch, M. Tarantola, E. Bodenschatz, W.-J. Rappel, Modeling self-organized spatio-temporal patterns of PIP3 and PTEN during spontaneous cell polarization. Phys Biol. 11, 046002 (2014).

62. D. Shao, W.-J. Rappel, H. Levine, Computational model for cell morphodynamics. Phys Rev Lett. 105, 108104 (2010).

63. B. A. Camley, Y. Zhao, B. Li, H. Levine, W.-J. Rappel, Crawling and turning in a minimal reaction-diffusion cell motility model: Coupling cell shape and biochemistry. Phys. Rev. E. 95, 012401–13 (2017).

64. E. Tunnacliffe, A. M. Corrigan, J. R. Chubb, Promoter-mediated diversification of transcriptional bursting dynamics following gene duplication. Proc Natl Acad Sci USA. 115, 8364–8369 (2018).

65. T. Gregor, K. Fujimoto, N. Masaki, S. Sawai, The onset of collective behavior in social amoebae. Science. 328, 1021–1025 (2010).

66. C. Okimura, A. Taniguchi, S. Nonaka, Y. Iwadate, Rotation of stress fibers as a single wheel in migrating fish keratocytes. Sci. Rep. 8, 1–10 (2018).

67. D. Taniguchi et al., Phase geometries of two-dimensional excitable waves govern self-organized morphodynamics of amoeboid cells. Proc Natl Acad Sci USA. 110, 5016–5021 (2013).

68. R. Winklbauer, Cell adhesion strength from cortical tension – an integration of concepts. J Cell Sci. 128, 3687–3693 (2015).

69. M. J. Footers, J. W. J. Kerssemakers, J. A. Theriot, M. Dogterom, Direct measurement of force generation by actin filament polymerization using an optical trap. Proc. Natl. Acad. Sci. USA. 104, 2181–2186 (2007).

70. M. P. Clausen, H. Colin-York, F. Schneider, C. Eggeling, M. Fritzsche, Dissecting the actin cortex density and membrane-cortex distance in living cells by super-resolution microscopy. J Phys D Appl Phys. 50, 064002 (2017).

71. S. Fukushima, S. Matsuoka, M. Ueda, Excitable dynamics of Ras triggers spontaneous symmetry breaking of PIP3 signaling in motile cells. J Cell Sci. 132, jcs224121 (2019).

72. A. T. Sasaki et al., G protein-independent Ras/PI3K/F-actin circuit regulates basic cell motility. J Cell Biol. 178, 185–191 (2007).

73. M.-J. Wang, Y. Artemenko, W.-J. Cai, P. A. Iglesias, P. N. Devreotes, The Directional Response of Chemotactic Cells Depends on a Balance between Cytoskeletal Architecture and the External Gradient. Cell Rep. 9, 1110–1121 (2014).

