## Supplementary Information for "Comparative mapping of crawling-cell morphodynamics in deep learning-based feature space"

### 1. Mathematical model

#### 1-1. Model equations

The reaction scheme for $U$ and $V$ describes two counteracting reactions U → V and V → U mediated by enzymes A and B. We adopted an earlier model (*67*) with a minor modification to include a feedback from an additional polarity factor W; namely,

$$\frac{\mathrm{d}A}{\mathrm{d}t} = k_1 W V A_{cyt} - k_2 A$$

$$\frac{\mathrm{d}B}{\mathrm{d}t} = k_3 U B_{cyt} - k_4 B$$

$$\frac{\mathrm{d}V}{\mathrm{d}t} = k_5 A U - k_6 B V - k_7 V$$

$$\frac{\mathrm{d}U}{\mathrm{d}t} = -k_5 A U + k_6 B V + k_8 - k_9 U$$

where $A$ and $B$ are the concentrations of enzymes in the membrane-bound form that mediate the reaction U → V and V → U, respectively. Originally, the forward reaction from $U$ to $V$ represented phosphorylation of the phosphatidylinositol (4,5)-bisphosphate (PIP2) to phosphatidylinositol (3,4,5)-triphosphate (PIP3) mediated by PI3kiase and the reverse reaction mediated by PTEN phosphatase, and the positive feedback amplification of $A$ described the F-actin dependent PI3K activation. Due to the introduction of new polarity variable $W$, the amplification rate for A in the present model assumed it to be proportional to $VW$ instead of $V^2$ (*67*). $A_{\mathrm{cyt}}$ and $B_{\mathrm{cyt}}$ represent the respective cytosolic fractions. Parameters $k_1$ through $k_9$ are the rate coefficients. We assume that $A_{\mathrm{cyt}}$ and $B_{\mathrm{cyt}}$ diffuse fast so that they are distributed uniformly in space and that the sum of membrane-bound and cytosolic fractions are conserved, $A_{\mathrm{cyt}}\int_\Omega \phi \mathrm{d}\boldsymbol{r} + \int_\Omega \phi A \mathrm{d}\boldsymbol{r} = A_{\mathrm{tot}}\int_\Omega \phi \mathrm{d}\boldsymbol{r}$ and $B_{\mathrm{cyt}}\int_\Omega \phi \mathrm{d}\boldsymbol{r} + \int_\Omega \phi B \mathrm{d}\boldsymbol{r} = B_{\mathrm{tot}}\int_\Omega \phi \mathrm{d}\boldsymbol{r}$ . Ω represents the entire spatial region computed. Provided that the reaction for $A$ and $B$ is fast compared to those of $U$ and $V$ (*67*), we obtain Equations (2a) and (2b) by combining the reactions with molecular diffusion in the phase-field $\phi$ (diffusion coefficients for $U$ and $V$ are $D_U$ and $D_V$) and the absorption of $U$ at the cell boundary represented by the last term of Eq.(2a). Parameters in Eqs. (2a) and (2b) are $\alpha = k_5A_{\mathrm{tot}}$, $\beta = k_6B_{\mathrm{tot}}$, $K_{\mathrm{k}} = k_2/k_1$, $K_{\mathrm{p}} = k_4/k_3$, $\mu = k_7$, $s = k_8$ and $\gamma = k_9$.

For the time evolution of $W$, we adopted an interconversion process between the active form $W$ and the inactive form $W$* under both positive and negative feedback regulation (fig. S3e).

$$\frac{\mathrm{d}X}{\mathrm{d}t} = \kappa_1 W^2 - \kappa_2 X \tag{S2-1}$$

$$\frac{\mathrm{d}Y}{\mathrm{d}t} = \kappa_3 + \kappa_4 W^2 - \kappa_5 Y \tag{S2-2}$$

$$\frac{\mathrm{d}W}{\mathrm{d}t} = \kappa_6 X W^* - \kappa_7 Y W \tag{S2-3}$$

where X mediates the reaction $W \to W^*$ and Y mediates $W^* \to W$. By assuming sufficiently fast relaxation for $X$ and $Y$ compared to $W$, we adiabatically eliminated the dynamics for $X$ and $Y$ ($\mathrm{d}X/\mathrm{d}t = 0$ and $\mathrm{d}Y/\mathrm{d}t = 0$) so that

$$\frac{\mathrm{d}W}{\mathrm{d}t} = k_{W1}(-\rho W^3 + \rho W^* W^2 - W)$$

(S2-4).

By further assuming that $W$ is positively regulated by $V$ following first-order kinetics and coupling the reactions with molecular diffusion in the phase-field $\phi$ (*62*), we obtained Eq.2-2c which is similar in form to the cell polarity model (*29*) derived from somewhat more detailed bistable regulation of Rho-GTPase. Other related studies (*38*, *63*) have also employed similar equations.

Under conditions where $W$ becomes enslaved to $V$ ($\rho \to 0$, $D_w \to 0$; Eq. 2c), the system is reduced to the 2-variable model (*67*) (Eq. 2a,b; fig. S3a-d). Similarly when activation of $W$ by $V$ is diminished ($\zeta \to 0$; Eq. 2c), the system becomes the 1-variable cell polarity model (fig. S3e-h). In the 2-variable regime, depending on the boundary conditions, excitable waves propagated either globally (*67*) or as randomly nucleated patches at the edge (fig. S3b) (see also (*61*) for a related excitable model). These protrusions were always localized in space and time and did not give rise to persistently polarized cell morphology. On the other hand, the 1-variable model exhibited polarized morphologies and new protrusions grew only in the rear and not in the anterior region (fig. S3g, h). Moreover, the average morphologies in the 1-variable regime (fig. S3i; small circles) were displaced towards higher PC1 from the *Dictyostelium* (agg) data (fig. S3i; oval regions) and thus mapped near the non-bifurcating ellipsoidal shapes analyzed in Fig. 1m.

**1-2. Numerical calculation**

Numerical calculation was performed by a semi-implicit type of explicit method (*63*). First, the updated values for $\phi$, $\phi V$, $\phi U$ and $\phi W$ are calculated as follows,

$$(\phi V)_{t+\Delta t} = \phi_t V_t + \Delta t \left(\frac{\partial(\phi V)}{\partial t}\right)$$

(S3-1),

$$(\phi U)_{t+\Delta t} = \phi_t U_t + \Delta t \left(\frac{\partial(\phi U)}{\partial t}\right)$$

(S3-2),

$$(\phi W)_{t+\Delta t} = \phi_t W_t + \Delta t \left(\frac{\partial(\phi W)}{\partial t}\right)$$

(S3-3),

$$\phi_{t+\Delta t} = \phi_t + \Delta t \left(\frac{\partial \phi}{\partial t}\right)$$

(S3-4).

Next, if $\phi_t$ is larger than $\mathrm{th}_\varphi$, $V$, $U$ and $W$ are updated following

$$V_{t+\Delta t} = \frac{(\phi V)_{t+\Delta t}}{\phi_{t+\Delta t}}$$

(S3-6),

$$U_{t+\Delta t} = \frac{(\phi U)_{t+\Delta t}}{\phi_{t+\Delta t}}$$

(S3-7),

$$W_{t+\Delta t} = \frac{(\phi W)_{t+\Delta t}}{\phi_{t+\Delta t}}$$

(S3-8),

otherwise

$$V_{t+\Delta t} = (\phi V)_{t+\Delta t}$$

(S3-9),

$$U_{t+\Delta t} = (\phi U)_{t+\Delta t}$$

(S3-10),

$$W_{t+\Delta t} = (\phi W)_{t+\Delta t}$$

(S3-11).

The value of $\mathrm{th}_\varphi$ was set to $10^{-4}$. The time increment ($\Delta t$) and the spatial grid ($\Delta x$) are set to $\Delta t = 4\times10^{-5}$ and $\Delta x = 0.1$. The program was coded in C++ with Open ACC (PGI compiler) for GPGPU computation (NVIDIA GeForce 1080 Ti). The noise terms in Eq. 2a and 2b were given by $N(\boldsymbol{x}, t) = N_0 \exp\left(-|\boldsymbol{x} - \boldsymbol{x}_c|^2/2d^2\right)$, where $\boldsymbol{x}_c$ is a random position selected at rate $\theta$ per area and $d$ is the nucleation size. The noise amplitude $N_0$ follows an exponential distribution with the average σ. In case $N(\boldsymbol{r},t)$ exceeds $U(\boldsymbol{r},t)$, $N(\boldsymbol{r},t)$ was reset to $U$ so as to prevent $U$ from taking negative values. Parameters were fixed to $\theta = 1.4$, $d = 0.8$, $\sigma = 0.075$ for all calculations on full 3-variable reaction regime. For 1-variable and 2-variable regime, σ was varied.

### 2. Feature extraction from the intermediate-layer of the trained neural network

To check robustness of the feature mapping, we revisited the morphology analysis using the 256-dimension intermediate-layer of the trained convolutional neural network (Fig. 1a). This was to check that important shape features useful to further constrain the model parameters were not lost in the final layer of the convolutional network. Starting from the 256 dimensional feature, we found that PCA yielded almost identical shape representation in two dimensional space. The contribution of the first two principle components PC1, PC2 and the sum of the all other components (PC3~PC256) were approximately 61.4%, 34.2% and 4.4%, respectively. Real dataset were clustered in the feature space accordingly to their respective attributes (fig. S6a). This new space was very similar to the one obtained from the last layer in terms of the relative positioning of the trained data. The vegetative *Dictyostelium* data are located between HL-60 and aggregation-stage *Dictyostelium,* and Nocodazole-treated HL-60 data are located between HL-60 and keratocyte, whereas *Dictyostelium racE*- data are mapped between aggregation-stage *Dictyostelium* and keratocyte. Mapping of simulated cell morphologies are shown in fig. S6b. Parameter sets were ranked according to the Euclidean distance in the 256 dimensional feature space between the simulated morphology and the mean features of the reference real data. In case of HL-60, keratocyte, nocodazole-treated HL-60 and *Dictyostelium racE*- data, top ranking simulations were identical to those obtained using the feature vector ***F*** described in the main text (Fig. 3f,i and Fig.5 e,g). The top ranking simulations for aggregation-stage *Dictyostelium* and vegetative *Dictyostelium* based on the 256-dimensional features (fig. S6c,d) were different (SI Table 9) but still within a very close range in the feature space (fig. S6b). These results indicate that the measure of similarity does not depend largely on whether the feature is based on the intermediate or the final layer of the trained network.

**Table S1. Data samples used for shape feature extraction (total snapshot images).**

| Dataset | ***Dictyostelium* (aggregation-stage)** | **HL-60** | **keratocyte** |
|---|---|---|---|
| Training | 2430 (1215) | 2286 (1143) | 2393 (731) |
| Validation | 296 (234) | 296 (296) | 296 (211) |

Sample # (Sample # before data augmentation)

**Table S2. Parameters for the phase-field dynamics (Eq. 1).**

| Parameter | Description | Value(s) | Misc. |
|---|---|---|---|
| $\tau$ | Viscous friction coefficient | $0.83 \times \tau'$ [pN/μm$^2$] | Fixed |
| $\eta$ | Surface tension | 1.0 [pN] | Fixed |
| $\varepsilon$ | Spatial scale of the phase boundary | 1.0 [μm] | Fixed |
| $M$ | Area conservation constraint | 0.5 [pN/μm$^3$] | Fixed |
| $A_0$ | Cell area | 78.83 [μm$^2$] | Fixed |
| $a_W$ | Protrusion force | 0.8, 1.6, 2.4, 3.2, 4.0 [pN/μm (conc.)$^{-1}$] | |

**Table S3: Parameters for the kinetic equations (Eqs. 2a-c).**

| Parameter | Description | Value(s) | Misc. |
|---|---|---|---|
| $\alpha$ | Relative strength of the feedback from V and W to reaction V → U | 2.0 | Fixed (for 1-variable regime, also $\alpha = 6, 10$) |
| $\beta$ | Relative strength of the feedback from U to reaction U → V | 3.5 | Fixed |
| $s$ | Supply rate of U | 1.0 | Fixed |
| $\gamma$ | Decay rate of U | 0.1, 0.3, 0.5, 0.7 | |
| $\mu$ | Decay rate of V | 0.1, 0.3, 0.5, 0.7, 0.9 | |
| $D_U$ | Diffusion coefficient of U | 0.05 | Fixed |
| $D_V$ | Diffusion coefficient of V | 0.2 | Fixed |
| $\chi_U$ | Absorption rate of *U* at the edge | 0, 10, 30, 50, 70 | |
| $\zeta$ | Activation rate of W by V | 200 | Fixed |
| $k_{W1}$ | Decay rate of W | 10, 20, 50, 90, 95, 100, 105, 110, 220 | |
| $\rho$ | Relative rate of W auto-regulation | 4.55, 4.76, 5, 5.26, 5.56 | |
| $D_W$ | Diffusion coefficient of W | 1.2, 3.0 | |
| $W_{tot}$ | Total amount of W and W* | 50, 60, 70, 80, 90, 100, 110, 120 | |
| $K_K$ | Effective Michaelis-Menten constant | 3.5 | Fixed |
| $K_P$ | Effective Michaelis-Menten constant | 3.2 | Fixed |
| $\tau'$ | Time scaling factor | 10 | |

**Table S4. Parameters for the 2-variable (*V*-*U*) equations.**

| $(\chi_{U}, \mu, \gamma, a_{V}, \alpha, \sigma)$ | | |
|---|---|---|
| 0, 0.1, 0.1, 24, 2, 0.075 | 0, 0.3, 0.1, 24, 2, 0.075 | 0, 0.1, 0.1, 8, 2, 0.3 |
| 0, 0.5, 0.1, 24, 2, 0.075 | 0, 0.7, 0.1, 24, 2, 0.075 | 0, 0.5, 0.1, 8, 2, 0.3 |
| 0, 0.5, 0.3, 24, 2, 0.075 | 0, 0.9, 0.1, 24, 2, 0.075 | 0, 0.5, 0.3, 8, 2, 0.3 |
| 0, 0.5, 0.5, 24, 2, 0.075 | 0, 0.5, 0.7, 24, 2, 0.075 | 0, 0.5, 0.5, 8, 2, 0.3 |
| 0, 0.1, 0.1, 24, 10, 0.075 | 0, 0.3, 0.1, 24, 10, 0.075 | 0, 0.3, 0.1, 8, 2, 0.3 |
| 0, 0.5, 0.1, 24, 10, 0.075 | 0, 0.7, 0.1, 24, 10, 0.075 | 0, 0.7, 0.1, 8, 2, 0.3 |
| 0, 0.5, 0.3, 24, 10, 0.075 | 0, 0.9, 0.1, 24, 10, 0.075 | 0, 0.9, 0.1, 8, 2, 0.3 |
| 0, 0.5, 0.5, 24, 10, 0.075 | 0, 0.5, 0.7, 24, 10, 0.075 | 0, 0.5, 0.7, 8, 2, 0.3 |
| 50, 0.1, 0.1, 24, 2, 0.075 | 50, 0.3, 0.1, 24, 2, 0.075 | 0, 0.1, 0.1, 8,10, 0.3 |
| 50, 0.5, 0.1, 24, 2, 0.075 | 50, 0.7, 0.1, 24, 2, 0.075 | 0, 0.5, 0.1, 8, 10, 0.3 |
| 50, 0.5, 0.3, 24, 2, 0.075 | 50, 0.9, 0.1, 24, 2, 0.075 | 0, 0.5, 0.3, 8, 10, 0.3 |
| 50, 0.5, 0.5, 24, 2, 0.075 | 50, 0.5, 0.7, 24, 2, 0.075 | 0, 0.5, 0.5, 8, 10, 0.3 |
| 50, 0.1, 0.1, 24, 10, 0.075 | 50, 0.3, 0.1, 24, 10, 0.075 | 0, 0.3, 0.1, 8, 10, 0.3 |
| 50, 0.5, 0.1, 24, 10, 0.075 | 50, 0.7, 0.1, 24, 10, 0.075 | 0, 0.7, 0.1, 8, 10, 0.3 |
| 50, 0.5, 0.3, 24, 10, 0.075 | 50, 0.9, 0.1, 24, 10, 0.075 | 0, 0.9, 0.1, 8, 10, 0.3 |
| 50, 0.5, 0.5, 24, 10, 0.075 | 50, 0.5, 0.7, 24, 10, 0.075 | 0, 0.5, 0.7, 8, 10, 0.3 |
| 0, 0.5, 0.5, 24, 6, 0.075 | 0, 0.3, 0.1, 24, 6, 0.075 | 50, 0.1, 0.1, 8, 2, 0.3 |
| 0, 0.1, 0.1, 24, 6, 0.075 | 0, 0.7, 0.1, 24, 6, 0.075 | 50, 0.5, 0.1, 8, 2, 0.3 |
| 0, 0.5, 0.1, 24, 6, 0.075 | 0, 0.9, 0.1, 24, 6, 0.075 | 50, 0.5, 0.3, 8, 2, 0.3 |
| 0, 0.5, 0.3, 24, 6, 0.075 | 0, 0.5, 0.7, 24, 6, 0.075 | 50, 0.5, 0.5, 8, 2, 0.3 |
| 50, 0.1, 0.1, 24, 6, 0.075 | 50, 0.3, 0.1, 24, 6, 0.075 | 50, 0.3, 0.1, 8, 2, 0.3 |
| 50, 0.5, 0.1, 24, 6, 0.075 | 50, 0.7, 0.1, 24, 6, 0.075 | 50, 0.7, 0.1, 8, 2, 0.3 |
| 50, 0.5, 0.3, 24, 6, 0.075 | 50, 0.9, 0.1, 24, 6, 0.075 | 50, 0.9, 0.1, 8, 2, 0.3 |
| 50, 0.5, 0.5, 24, 6, 0.075 | 50, 0.5, 0.7, 24, 6, 0.075 | 50, 0.5, 0.7, 8, 2, 0.3 |

**Table S5. Parameters for the 1-variable (*W*) equation.**

| $(k_{W1}, \rho, a_{W}, D_{W}, W_{tot}, \sigma)$ | | |
|---|---|---|
| 10, 5, 2.4, 3, 80, 0.15 | 110, 4.5455, 2.4, 3, 80, 0.5 | 75, 6.6667, 1.6, 3, 80, 0.15 |
| 20, 5, 2.4, 3, 80, 0.15 | 110, 4.5455, 0.8, 3, 80, 0.5 | 85, 5.8824, 1.6, 3, 80, 0.15 |
| 50, 5, 2.4, 3, 80, 0.15 | 110, 4.5455, 1.6, 3,80, 0.5 | 90, 5.5556, 1.6, 3, 100, 0.15 |
| 100, 5, 2.4, 3, 80, 0.15 | 110, 4.5455, 3.2, 3,80, 0.5 | 90, 5.5556, 0.8, 3, 100, 0.15 |
| 90, 5.5556, 2.4, 3, 80, 0.15 | 110, 4.5455, 4,3, 80, 0.5 | 90, 7.6923, 1.6, 3, 80, 0.15 |
| 95, 5.2632, 2.4, 3, 80, 0.15 | 110, 4.5455, 2.4, 3, 60, 0.5 | 45, 5.5556, 1.6, 3, 100, 0.15 |
| 105, 4.7619, 2.4, 3, 80, 0.15 | 110, 4.5455, 2.4, 3, 70, 0.5 | 90, 5.5556, 0.4, 3, 100, 0.15 |
| 110, 4.5455, 2.4, 3, 80, 0.15 | 110, 4.5455, 2.4, 3, 90, 0.5 | 45, 5.5556, 0.8, 3, 100, 0.15 |
| 110, 4.5455, 0.8, 3, 80, 0.15 | 110, 4.5455, 2.4, 3, 100, 0.5 | 70, 7.1429, 0.4, 3, 80, 0.15 |
| 110, 4.5455, 1.6, 3, 80, 0.15 | 110, 4.5455, 2.4, 3, 110, 0.5 | 35, 7.1429, 0.8, 3, 80, 0.15 |
| 110, 4.5455, 3.2, 3, 80, 0.15 | 110, 4.5455, 2.4, 3, 110, 0.15 | 70, 7.1429, 0.4, 3, 100, 0.15 |
| 110, 4.5455, 4, 3, 80, 0.15 | 110, 4.5455, 2.4, 3, 120, 0.5 | 80, 6.2500, 0.4, 3, 100, 0.15 |
| 110, 4.5455, 2.4, 3, 60, 0.15 | 110, 4.5455, 2.4, 3, 120, 0.15 | 35, 7.1429, 0.8, 3, 100, 0.15 |
| 110, 4.5455, 2.4, 3, 70, 0.15 | 80, 6.25, 0.8, 3, 80, 0.15 | 40, 6.2500, 0.8, 3, 100, 0.15 |
| 110, 4.5455, 2.4, 3, 90, 0.15 | 80, 6.25, 1.6, 3, 80, 0.15 | 80, 6.2500, 0.4, 3, 90, 0.15 |
| 110, 4.5455, 2.4, 3, 100, 0.15 | 80, 6.25, 3.2, 3, 80, 0.15 | 40, 6.2500, 0.8, 3, 90, 0.15 |
| 90, 5.5556, 2.4, 3, 80, 0.5 | 80, 6.25, 4, 3, 80, 0.15 | 80, 6.2500, 0.8, 3, 90, 0.15 |
| 95, 5.2632, 2.4, 3, 80, 0.5 | 90, 5.5556, 1.6, 3, 80, 0.15 | 40, 6.2500, 1.6, 3, 90, 0.15 |
| 105, 4.7619, 2.4, 3, 80, 0.5 | 100, 5, 1.6, 3, 80, 0.15 | |

**Table S6. Model parameters (228 sets).**

| ($\chi_U$, $k_{W1}$, $\mu$, $\rho$, $\gamma$, $a_W$, $D_W$, $W_{tot}$) | | |
|---|---|---|
| 0,110,0.5,4.5455,0.1,0.8,1.2,80 | 50,110,0.5,4.5455,0.5,1.6,1.2,80 | 0,200,0.5,5,0.1,2.4,3,80 |
| 0,110,0.5,4.5455,0.1,1.6,1.2,80 | 50,110,0.5,4.5455,0.5,2.4,1.2,80 | 0,200,0.7,5,0.1,2.4,3,80 |
| 0,110,0.5,4.5455,0.1,2.4,1.2,80 | 50,110,0.5,4.5455,0.5,3.2,1.2,80 | 0,200,0.9,5,0.1,2.4,3,80 |
| 0,110,0.5,4.5455,0.1,3.2,1.2,80 | 50,110,0.5,4.5455,0.5,4,1.2,80 | 0,90,0.5,5.5556,0.1,2.4,3,80 |
| 0,110,0.5,4.5455,0.1,4,1.2,80 | 50,110,0.5,4.5455,0.7,0.8,1.2,80 | 0,95,0.5,5.2632,0.1,2.4,3,80 |
| 0,110,0.5,4.5455,0.3,0.8,1.2,80 | 50,110,0.5,4.5455,0.7,1.6,1.2,80 | 0,100,0.5,5,0.1,2.4,3,80 |
| 0,110,0.5,4.5455,0.3,1.6,1.2,80 | 50,110,0.5,4.5455,0.7,2.4,1.2,80 | 0,105,0.5,4.7619,0.1,2.4,3,80 |
| 0,110,0.5,4.5455,0.3,2.4,1.2,80 | 50,110,0.5,4.5455,0.7,3.2,1.2,80 | 0,110,0.5,4.5455,0.1,2.4,3,80 |
| 0,110,0.5,4.5455,0.3,3.2,1.2,80 | 50,110,0.5,4.5455,0.7,4,1.2,80 | 0,90,0.5,5.5556,0.3,2.4,3,80 |
| 0,110,0.5,4.5455,0.3,4,1.2,80 | 0,110,0.5,4.5455,0.1,2.4,1.2,50 | 0,95,0.5,5.2632,0.3,2.4,3,80 |
| 0,110,0.5,4.5455,0.5,0.8,1.2,80 | 0,110,0.5,4.5455,0.1,2.4,1.8,50 | 0,100,0.5,5,0.3,2.4,3,80 |
| 0,110,0.5,4.5455,0.5,1.6,1.2,80 | 0,110,0.5,4.5455,0.1,2.4,2.4,50 | 0,105,0.5,4.7619,0.3,2.4,3,80 |
| 0,110,0.5,4.5455,0.5,2.4,1.2,80 | 50,110,0.5,4.5455,0.1,2.4,3,50 | 0,110,0.5,4.5455,0.3,2.4,3,80 |
| 0,110,0.5,4.5455,0.5,3.2,1.2,80 | 50,110,0.5,4.5455,0.1,2.4,1.2,60 | 0,90,0.5,5.5556,0.5,2.4,3,80 |
| 0,110,0.5,4.5455,0.5,4,1.2,80 | 0,110,0.5,4.5455,0.1,2.4,1.8,60 | 0,95,0.5,5.2632,0.5,2.4,3,80 |
| 0,110,0.5,4.5455,0.7,0.8,1.2,80 | 0,110,0.5,4.5455,0.1,2.4,2.4,60 | 0,100,0.5,5,0.5,2.4,3,80 |
| 0,110,0.5,4.5455,0.7,1.6,1.2,80 | 50,110,0.5,4.5455,0.1,2.4,3,60 | 0,105,0.5,4.7619,0.5,2.4,3,80 |
| 0,110,0.5,4.5455,0.7,2.4,1.2,80 | 50,110,0.5,4.5455,0.1,2.4,1.2,70 | 0,110,0.5,4.5455,0.5,2.4,3,80 |
| 0,110,0.5,4.5455,0.7,3.2,1.2,80 | 0,110,0.5,4.5455,0.1,2.4,1.8,70 | 0,90,0.5,5.5556,0.7,2.4,3,80 |
| 0,110,0.5,4.5455,0.7,4,1.2,80 | 0,110,0.5,4.5455,0.1,2.4,2.4,70 | 0,95,0.5,5.2632,0.7,2.4,3,80 |
| 0,110,0.5,4.5455,0.1,2.4,1.2,50 | 50,110,0.5,4.5455,0.1,2.4,3,70 | 0,100,0.5,5,0.7,2.4,3,80 |
| 0,110,0.5,4.5455,0.1,2.4,1.8,50 | 0,110,0.5,4.5455,0.1,2.4,1.2,80 | 0,105,0.5,4.7619,0.7,2.4,3,80 |
| 0,110,0.5,4.5455,0.1,2.4,2.4,50 | 0,110,0.5,4.5455,0.1,2.4,1.8,80 | 0,110,0.5,4.5455,0.7,2.4,3,80 |
| 0,110,0.5,4.5455,0.1,2.4,3,50 | 0,110,0.5,4.5455,0.1,2.4,2.4,80 | 0,90,0.5,5.5556,0.1,2.4,3,80 |
| 0,110,0.5,4.5455,0.1,2.4,1.2,60 | 50,110,0.5,4.5455,0.1,2.4,3,80 | 0,95,0.5,5.2632,0.1,2.4,3,80 |
| 0,110,0.5,4.5455,0.1,2.4,1.8,60 | 50,110,0.5,4.5455,0.1,2.4,1.2,90 | 0,100,0.5,5,0.1,2.4,3,80 |
| 0,110,0.5,4.5455,0.1,2.4,2.4,60 | 0,110,0.5,4.5455,0.1,2.4,1.8,90 | 0,105,0.5,4.7619,0.1,2.4,3,80 |
| 0,110,0.5,4.5455,0.1,2.4,3,60 | 0,110,0.5,4.5455,0.1,2.4,2.4,90 | 0,110,0.5,4.5455,0.1,2.4,3,80 |
| 0,110,0.5,4.5455,0.1,2.4,1.2,70 | 50,110,0.5,4.5455,0.1,2.4,3,90 | 0,90,0.5,5.5556,0.3,2.4,3,80 |
| 0,110,0.5,4.5455,0.1,2.4,1.8,70 | 50,110,0.5,4.5455,0.1,2.4,1.2,100 | 0,95,0.5,5.2632,0.3,2.4,3,80 |
| 0,110,0.5,4.5455,0.1,2.4,2.4,70 | 0,110,0.5,4.5455,0.1,2.4,1.8,100 | 0,100,0.5,5,0.3,2.4,3,80 |
| 0,110,0.5,4.5455,0.1,2.4,3,70 | 0,110,0.5,4.5455,0.1,2.4,2.4,100 | 0,105,0.5,4.7619,0.3,2.4,3,80 |
| 0,110,0.5,4.5455,0.1,2.4,1.2,80 | 50,110,0.5,4.5455,0.1,2.4,3,100 | 0,110,0.5,4.5455,0.3,2.4,3,80 |
| 0,110,0.5,4.5455,0.1,2.4,1.8,80 | 50,110,0.5,4.5455,0.1,2.4,1.2,110 | 0,90,0.5,5.5556,0.5,2.4,3,80 |
| 0,110,0.5,4.5455,0.1,2.4,2.4,80 | 0,110,0.5,4.5455,0.1,2.4,1.8,110 | 0,95,0.5,5.2632,0.5,2.4,3,80 |
| 0,110,0.5,4.5455,0.1,2.4,3,80 | 0,110,0.5,4.5455,0.1,2.4,2.4,110 | 0,100,0.5,5,0.5,2.4,3,80 |
| 0,110,0.5,4.5455,0.1,2.4,1.2,90 | 50,110,0.5,4.5455,0.1,2.4,3,110 | 0,105,0.5,4.7619,0.5,2.4,3,80 |
| 0,110,0.5,4.5455,0.1,2.4,1.8,90 | 0,10,0.1,5,0.1,2.4,3,80 | 0,110,0.5,4.5455,0.5,2.4,3,80 |
| 0,110,0.5,4.5455,0.1,2.4,2.4,90 | 0,10,0.3,5,0.1,2.4,3,80 | 0,90,0.5,5.5556,0.7,2.4,3,80 |
| 0,110,0.5,4.5455,0.1,2.4,3,90 | 0,10,0.5,5,0.1,2.4,3,80 | 0,95,0.5,5.2632,0.7,2.4,3,80 |
| 0,110,0.5,4.5455,0.1,2.4,1.2,100 | 0,10,0.7,5,0.1,2.4,3,80 | 0,100,0.5,5,0.7,2.4,3,80 |
| 0,110,0.5,4.5455,0.1,2.4,1.8,100 | 0,10,0.9,5,0.1,2.4,3,80 | 0,105,0.5,4.7619,0.7,2.4,3,80 |
| 0,110,0.5,4.5455,0.1,2.4,2.4,100 | 0,20,0.1,5,0.1,2.4,3,80 | 0,110,0.5,4.5455,0.7,2.4,3,80 |
| 0,110,0.5,4.5455,0.1,2.4,3,100 | 0,20,0.3,5,0.1,2.4,3,80 | 50,10,0.1,5,0.1,2.4,3,80 |
| 0,110,0.5,4.5455,0.1,2.4,1.2,110 | 0,20,0.5,5,0.1,2.4,3,80 | 50,10,0.3,5,0.1,2.4,3,80 |
| 0,110,0.5,4.5455,0.1,2.4,1.8,110 | 0,20,0.7,5,0.1,2.4,3,80 | 50,10,0.5,5,0.1,2.4,3,80 |
| 0,110,0.5,4.5455,0.1,2.4,2.4,110 | 0,20,0.9,5,0.1,2.4,3,80 | 50,10,0.7,5,0.1,2.4,3,80 |
| 0,110,0.5,4.5455,0.1,2.4,3,110 | 0,50,0.1,5,0.1,2.4,3,80 | 50,10,0.9,5,0.1,2.4,3,80 |
| 50,110,0.5,4.5455,0.1,0.8,1.2,80 | 0,50,0.3,5,0.1,2.4,3,80 | 50,20,0.1,5,0.1,2.4,3,80 |
| 50,110,0.5,4.5455,0.1,1.6,1.2,80 | 0,50,0.5,5,0.1,2.4,3,80 | 50,20,0.3,5,0.1,2.4,3,80 |
| 50,110,0.5,4.5455,0.1,2.4,1.2,80 | 0,50,0.7,5,0.1,2.4,3,80 | 50,20,0.5,5,0.1,2.4,3,80 |
| 50,110,0.5,4.5455,0.1,3.2,1.2,80 | 0,50,0.9,5,0.1,2.4,3,80 | 50,20,0.7,5,0.1,2.4,3,80 |
| 50,110,0.5,4.5455,0.1,4,1.2,80 | 0,100,0.1,5,0.1,2.4,3,80 | 50,20,0.9,5,0.1,2.4,3,80 |
| 50,110,0.5,4.5455,0.3,0.8,1.2,80 | 0,100,0.3,5,0.1,2.4,3,80 | 50,50,0.1,5,0.1,2.4,3,80 |
| 50,110,0.5,4.5455,0.3,1.6,1.2,80 | 0,100,0.5,5,0.1,2.4,3,80 | 50,50,0.3,5,0.1,2.4,3,80 |
| 50,110,0.5,4.5455,0.3,2.4,1.2,80 | 0,100,0.7,5,0.1,2.4,3,80 | 50,50,0.5,5,0.1,2.4,3,80 |
| 50,110,0.5,4.5455,0.3,3.2,1.2,80 | 0,100,0.9,5,0.1,2.4,3,80 | 50,50,0.5,5,0.1,2.4,3,80 |
| 50,110,0.5,4.5455,0.3,4,1.2,80 | 0,200,0.1,5,0.1,2.4,3,80 | 50,50,0.9,5,0.1,2.4,3,80 |
| 50,110,0.5,4.5455,0.5,0.8,1.2,80 | 0,200,0.3,5,0.1,2.4,3,80 | 50,100,0.1,5,0.1,2.4,3,80 |

| |
|---|
| 50,100,0.3,5,0.1,2.4,3,80<br>50,100,0.5,5,0.1,2.4,3,80<br>50,100,0.7,5,0.1,2.4,3,80<br>50,100,0.9,5,0.1,2.4,3,80<br>50,200,0.1,5,0.1,2.4,3,80<br>50,200,0.3,5,0.1,2.4,3,80<br>50,200,0.5,5,0.1,2.4,3,80<br>50,200,0.7,5,0.1,2.4,3,80<br>50,200,0.9,5,0.1,2.4,3,80<br>50,90,0.5,5.5556,0.1,2.4,3,80<br>50,95,0.5,5.2632,0.1,2.4,3,80<br>50,100,0.5,5,0.1,2.4,3,80<br>50,105,0.5,4.7619,0.1,2.4,3,80<br>50,110,0.5,4.5455,0.1,2.4,3,80<br>50,90,0.5,5.5556,0.3,2.4,3,80<br>50,95,0.5,5.2632,0.3,2.4,3,80<br>50,100,0.5,5,0.3,2.4,3,80<br>50,105,0.5,4.7619,0.3,2.4,3,80<br>50,110,0.5,4.5455,0.3,2.4,3,80<br>50,90,0.5,5.5556,0.5,2.4,3,80<br>50,95,0.5,5.2632,0.5,2.4,3,80<br>50,100,0.5,5,0.5,2.4,3,80<br>50,105,0.5,4.7619,0.5,2.4,3,80<br>50,110,0.5,4.5455,0.5,2.4,3,80<br>50,90,0.5,5.5556,0.7,2.4,3,80<br>50,95,0.5,5.2632,0.7,2.4,3,80<br>50,100,0.5,5,0.7,2.4,3,80<br>50,105,0.5,4.7619,0.7,2.4,3,80<br>50,110,0.5,4.5455,0.7,2.4,3,80<br>50,90,0.5,5.5556,0.1,2.4,3,80<br>50,95,0.5,5.2632,0.1,2.4,3,80<br>50,100,0.5,5,0.1,2.4,3,80<br>50,105,0.5,4.7619,0.1,2.4,3,80<br>50,110,0.5,4.5455,0.1,2.4,3,80<br>50,90,0.5,5.5556,0.3,2.4,3,80<br>50,95,0.5,5.2632,0.3,2.4,3,80<br>50,100,0.5,5,0.3,2.4,3,80<br>50,105,0.5,4.7619,0.3,2.4,3,80<br>50,110,0.5,4.5455,0.3,2.4,3,80<br>50,90,0.5,5.5556,0.5,2.4,3,80<br>50,95,0.5,5.2632,0.5,2.4,3,80<br>50,100,0.5,5,0.5,2.4,3,80<br>50,105,0.5,4.7619,0.5,2.4,3,80<br>50,110,0.5,4.5455,0.5,2.4,3,80<br>50,90,0.5,5.5556,0.7,2.4,3,80<br>50,95,0.5,5.2632,0.7,2.4,3,80<br>50,100,0.5,5,0.7,2.4,3,80<br>50,105,0.5,4.7619,0.7,2.4,3,80<br>50,110,0.5,4.5455,0.7,2.4,3,80 |

**Table S7 Top ranking parameters for Score-D, Score-H and Score-K.**

| Reference morphology | $\|\boldsymbol{F}\|_2$ | CNN layer | score rank | $\chi_U$ | $k_{W1}$ | $\mu$ | $\rho$ | $\gamma$ | $a_W$ | $D_W$ | $W_{tot}$ |
|---|---|---|---|---|---|---|---|---|---|---|---|
| Score-D (agg) | 4954 | Final layer | 1 | 0 | 110.0 | 0.5 | 4.5455 | 0.3 | 4.0 | 1.2 | 80.0 |
| | 11637 | | 2 | 0 | 110.0 | 0.5 | 4.5455 | 0.7 | 4.0 | 1.2 | 80.0 |
| | 12855 | | 3 | 50.0 | 110.0 | 0.5 | 4.5455 | 0.5 | 4.0 | 1.2 | 80.0 |
| | 13917 | | 4 | 0 | 110.0 | 0.5 | 4.5455 | 0.5 | 4.0 | 1.2 | 80.0 |
| | 15273 | | 5 | 50.0 | 110.0 | 0.5 | 4.5455 | 0.7 | 4.0 | 1.2 | 80.0 |
| | 17338 | | 6 | 0 | 110.0 | 0.5 | 4.5455 | 0.5 | 3.2 | 1.2 | 80.0 |
| | 18571 | | 7 | 50.0 | 110.0 | 0.5 | 4.5455 | 0.3 | 4.0 | 1.2 | 80.0 |
| | 18933 | | 8 | 50.0 | 110.0 | 0.5 | 4.5455 | 0.1 | 4.0 | 1.2 | 80.0 |
| | 19327 | | 9 | 0 | 110.0 | 0.5 | 4.5455 | 0.1 | 4.0 | 1.2 | 80.0 |
| | 20461 | | 10 | 50.0 | 110.0 | 0.5 | 4.5455 | 0.5 | 3.2 | 1.2 | 80.0 |
| | 20570 | | 11 | 0 | 110.0 | 0.5 | 4.5455 | 0.3 | 2.4 | 3.0 | 80.0 |
| Score-H | 2561 | Final layer | 1 | 50.0 | 100.0 | 0.5 | 5.0 | 0.1 | 1.6 | 3.0 | 80.0 |
| | 3350 | | 2 | 0 | 90.0 | 0.5 | 5.5556 | 0.3 | 2.4 | 3.0 | 80.0 |
| | 4447 | | 3 | 0 | 90.0 | 0.5 | 5.5556 | 0.1 | 2.4 | 3.0 | 80.0 |
| | 5913 | | 4 | 0 | 95.0 | 0.5 | 5.2632 | 0.1 | 2.4 | 3.0 | 80.0 |
| | 6320 | | 5 | 0 | 100.0 | 0.1 | 5.0 | 0.1 | 2.4 | 3.0 | 80.0 |
| | 6420 | | 6 | 50.0 | 100.0 | 0.7 | 5.0 | 0.1 | 2.4 | 3.0 | 80.0 |
| | 6564 | | 7 | 0 | 50.0 | 0.3 | 5.0 | 0.1 | 2.4 | 3.0 | 80.0 |
| | 7235 | | 8 | 50.0 | 110.0 | 0.5 | 4.5455 | 0.1 | 1.6 | 1.2 | 80.0 |
| | 7804 | | 9 | 0 | 50.0 | 0.7 | 5.0 | 0.1 | 2.4 | 3.0 | 80.0 |
| | 8075 | | 10 | 0 | 100.0 | 0.5 | 5.0 | 0.1 | 1.6 | 3.0 | 80.0 |
| | 8405 | | 11 | 50.0 | 110.0 | 0.5 | 4.5455 | 0.1 | 0.8 | 1.2 | 80.0 |
| Score-K | 18021 | Final layer | 1 | 50.0 | 20.0 | 0.1 | 5.0 | 0.1 | 2.4 | 3.0 | 80.0 |
| | 18189 | | 2 | 50.0 | 50.0 | 0.1 | 5.0 | 0.1 | 2.4 | 3.0 | 80.0 |
| | 26460 | | 3 | 50.0 | 10.0 | 0.1 | 5.0 | 0.1 | 2.4 | 3.0 | 80.0 |
| | 40680 | | 4 | 50.0 | 110.0 | 0.5 | 4.5455 | 0.1 | 2.4 | 1.2 | 100.0 |
| | 41350 | | 5 | 50.0 | 100.0 | 0.5 | 5.0 | 0.1 | 1.6 | 3.0 | 100.0 |
| | 44566 | | 6 | 50.0 | 100.0 | 0.1 | 5.0 | 0.1 | 2.4 | 3.0 | 80.0 |
| | 50622 | | 7 | 50.0 | 110.0 | 0.5 | 4.5455 | 0.1 | 2.4 | 1.2 | 90.0 |
| | 55830 | | 8 | 50.0 | 20.0 | 0.3 | 5.0 | 0.1 | 2.4 | 3.0 | 80.0 |
| | 56589 | | 9 | 0 | 110.0 | 0.5 | 4.5455 | 0.1 | 2.4 | 1.8 | 100.0 |
| | 59338 | | 10 | 0 | 110.0 | 0.5 | 4.5455 | 0.1 | 2.4 | 1.2 | 100.0 |
| | 60322 | | 11 | 50.0 | 110.0 | 0.5 | 4.5455 | 0.1 | 2.4 | 1.8 | 100.0 |

**Table S8 Top ranking parameters for Score-D (veg) and Score-H (nocodazole) and Score-D (*racE*-) .**

| Reference morphology | $\|\|\boldsymbol{F}\|\|_2$ | CNN layer | score rank | $\chi_U$ | $k_{W1}$ | $\mu$ | $\rho$ | $\gamma$ | $a_W$ | $D_W$ | $W_{tot}$ |
|---|---|---|---|---|---|---|---|---|---|---|---|
| Score-D (veg) | 5671 | Final layer | 1 | 0 | 50.0 | 0.9 | 5.0 | 0.1 | 2.4 | 3.0 | 80.0 |
| | 5798 | | 2 | 50.0 | 110.0 | 0.5 | 4.5455 | 0.1 | 2.4 | 3.0 | 80.0 |
| | 6238 | | 3 | 50.0 | 100.0 | 0.9 | 5.0 | 0.1 | 2.4 | 3.0 | 80.0 |
| | 6382 | | 4 | 0 | 200.0 | 0.3 | 5.0 | 0.1 | 2.4 | 3.0 | 80.0 |
| | 6434 | | 5 | 0 | 100.0 | 0.7 | 5.0 | 0.1 | 2.4 | 3.0 | 80.0 |
| | 6742 | | 6 | 0 | 100.0 | 0.9 | 5.0 | 0.1 | 2.4 | 3.0 | 80.0 |
| | 7058 | | 7 | 50.0 | 100.0 | 0.5 | 5.0 | 0.1 | 2.4 | 3.0 | 80.0 |
| | 7239 | | 8 | 50.0 | 200.0 | 0.5 | 5.0 | 0.1 | 2.4 | 3.0 | 80.0 |
| | 7277 | | 9 | 50.0 | 200.0 | 0.3 | 5.0 | 0.1 | 2.4 | 3.0 | 80.0 |
| | 7495 | | 10 | 50.0 | 100.0 | 0.5 | 5.0 | 0.3 | 2.4 | 3.0 | 80.0 |
| | 7509 | | 11 | 50.0 | 200.0 | 0.7 | 5.0 | 0.1 | 2.4 | 3.0 | 80.0 |
| (agg) to (veg) comparison | | Final layer | | - | n/a | - | n/a | ** | ** | n/a | - |
| Score-H (nocodazole) | 2568 | Final layer | 1 | 0 | 100.0 | 0.5 | 5.0 | 0.1 | 1.6 | 3.0 | 90.0 |
| | 5559 | | 2 | 50.0 | 10.0 | 0.3 | 5.0 | 0.1 | 2.4 | 3.0 | 80.0 |
| | 5668 | | 3 | 0 | 100.0 | 0.5 | 5.0 | 0.1 | 2.4 | 3.0 | 90.0 |
| | 5906 | | 4 | 0 | 110.0 | 0.5 | 4.5455 | 0.1 | 2.4 | 3.0 | 110.0 |
| | 6821 | | 5 | 0 | 10.0 | 0.3 | 5.0 | 0.1 | 2.4 | 3.0 | 80.0 |
| | 7916 | | 6 | 0 | 110.0 | 0.5 | 4.5455 | 0.1 | 2.4 | 1.8 | 110.0 |
| | 9075 | | 7 | 50.0 | 20.0 | 0.5 | 5.0 | 0.1 | 2.4 | 3.0 | 80.0 |
| | 9402 | | 8 | 50.0 | 100.0 | 0.5 | 5.0 | 0.1 | 0.8 | 3.0 | 90.0 |
| | 11260 | | 9 | 50.0 | 20.0 | 0.9 | 5.0 | 0.1 | 2.4 | 3.0 | 80.0 |
| | 11278 | | 10 | 0 | 110.0 | 0.5 | 4.5455 | 0.1 | 2.4 | 1.2 | 110.0 |
| | 11316 | | 11 | 0 | 100.0 | 0.5 | 5.0 | 0.1 | 1.6 | 3.0 | 100.0 |
| Score-D (*racE*-) | 5664 | Final layer | 1 | 50.0 | 110.0 | 0.5 | 4.5455 | 0.1 | 2.4 | 1.2 | 90.0 |
| | 12327 | | 2 | 50.0 | 110.0 | 0.5 | 4.5455 | 0.1 | 2.4 | 1.2 | 100.0 |
| | 17115 | | 3 | 50.0 | 110.0 | 0.5 | 4.5455 | 0.1 | 2.4 | 1.8 | 100.0 |
| | 20419 | | 4 | 50.0 | 110.0 | 0.5 | 4.5455 | 0.1 | 2.4 | 2.4 | 100.0 |
| | 22708 | | 5 | 50.0 | 100.0 | 0.5 | 5.0 | 0.1 | 1.6 | 3.0 | 100.0 |
| | 24674 | | 6 | 50.0 | 110.0 | 0.5 | 4.5455 | 0.1 | 2.4 | 3.0 | 100.0 |
| | 26676 | | 7 | 50.0 | 10.0 | 0.1 | 5.0 | 0.1 | 2.4 | 3.0 | 80.0 |
| | 28313 | | 8 | 0 | 110.0 | 0.5 | 4.5455 | 0.1 | 2.4 | 1.8 | 100.0 |
| | 28548 | | 9 | 50.0 | 100.0 | 0.5 | 5.0 | 0.1 | 2.4 | 3.0 | 90.0 |
| | 28812 | | 10 | 50.0 | 100.0 | 0.5 | 5.0 | 0.1 | 2.4 | 3.0 | 100.0 |
| | 30237 | | 11 | 0 | 100.0 | 0.5 | 5.0 | 0.1 | 2.4 | 3.0 | 100.0 |
| (*racE*-) to (agg) comparison | | Final layer | | - | n/a | - | n/a | ** | ** | n/a | ** |
| (*racE*-) to (veg) comparison | | Final layer | | - | | * | n/a | - | - | n/a | ** |

*… ($p$ < 10%), **… ($p$ < 0.1%), ***… ($p$ < 0.001%), -… not significant

n/a : subject to bias from the choice of parameter search

**Table S9. Top ranking parameters for Score-D (veg) and Score-D (agg) based on feature extraction at the intermediate layer .**

| Reference morphology | $\|\|\boldsymbol{F}\|\|_2$ | CNN layer | rank | $\chi_U$ | $k_{W1}$ | $\mu$ | $\rho$ | $\gamma$ | $a_W$ | $D_W$ | $W_{tot}$ | Table S8 rank |
|---|---|---|---|---|---|---|---|---|---|---|---|---|
| Score-D (veg) | 2701 | Intermediate layer | 1 | 0.0 | 90.0 | 0.5 | 5.5556 | 0.5 | 2.4 | 3.0 | 80.0 | 21 |
| | 2744 | | 2 | 50.0 | 100.0 | 0.5 | 5.0 | 0.3 | 2.4 | 3.0 | 80.0 | 10 |
| | 2880 | | 3 | 50.0 | 105.0 | 0.5 | 4.7619 | 0.1 | 2.4 | 3.0 | 80.0 | 12 |
| | 2988 | | 4 | 50.0 | 100.0 | 0.5 | 5.0 | 0.1 | 2.4 | 3.0 | 80.0 | 7 |
| | 3083 | | 5 | 50.0 | 95.0 | 0.5 | 5.2632 | 0.5 | 2.4 | 3.0 | 80.0 | 30 |
| | 3190 | | 6 | 0.0 | 50.0 | 0.5 | 5.0 | 0.1 | 2.4 | 3.0 | 80.0 | 13 |
| | 3375 | | 7 | 50.0 | 110.0 | 0.5 | 4.5455 | 0.1 | 2.4 | 1.2 | 80.0 | 20 |
| | 3378 | | 8 | 50.0 | 200.0 | 0.3 | 5.0 | 0.1 | 2.4 | 3.0 | 80.0 | 9 |
| | 3382 | | 9 | 50.0 | 100.0 | 0.5 | 5.0 | 0.5 | 2.4 | 3.0 | 80.0 | 32 |
| | 3393 | | 10 | 50.0 | 110.0 | 0.5 | 4.5455 | 0.3 | 1.6 | 1.2 | 80.0 | 39 |
| | 3433 | | 11 | 50.0 | 95.0 | 0.5 | 5.2632 | 0.1 | 2.4 | 3.0 | 80.0 | 47 |
| Score-D (agg) | 2634 | Intermediate layer | 1 | 0.0 | 110.0 | 0.5 | 4.5455 | 0.7 | 4.0 | 1.2 | 80.0 | 2 |
| | 2707 | | 2 | 50.0 | 110.0 | 0.5 | 4.5455 | 0.5 | 4.0 | 1.2 | 80.0 | 3 |
| | 2972 | | 3 | 0.0 | 110.0 | 0.5 | 4.5455 | 0.3 | 4.0 | 1.2 | 80.0 | 1 |
| | 3123 | | 4 | 0.0 | 110.0 | 0.5 | 4.5455 | 0.5 | 4.0 | 1.2 | 80.0 | 4 |
| | 3276 | | 5 | 50.0 | 110.0 | 0.5 | 4.5455 | 0.7 | 4.0 | 1.2 | 80.0 | 5 |
| | 3969 | | 6 | 50.0 | 110.0 | 0.5 | 4.5455 | 0.3 | 4.0 | 1.2 | 80.0 | 7 |
| | 4153 | | 7 | 50.0 | 110.0 | 0.5 | 4.5455 | 0.1 | 4.0 | 1.2 | 80.0 | 8 |
| | 4268 | | 8 | 0.0 | 110.0 | 0.5 | 4.5455 | 0.5 | 3.2 | 1.2 | 80.0 | 6 |
| | 4547 | | 9 | 50.0 | 110.0 | 0.5 | 4.5455 | 0.5 | 3.2 | 1.2 | 80.0 | 10 |
| | 4588 | | 10 | 0.0 | 110.0 | 0.5 | 4.5455 | 0.1 | 4.0 | 1.2 | 80.0 | 9 |
| | 4885 | | 11 | 50.0 | 110.0 | 0.5 | 4.5455 | 0.3 | 3.2 | 1.2 | 80.0 | 13 |
| (agg) to (veg) comparison | | Intermediate layer | | - | n/a | - | n/a | * | *** | n/a | - | |
| Score-D (*racE*-) | 1719 | Intermediate layer | 1 | 50.0 | 110.0 | 0.5 | 4.5455 | 0.1 | 2.4 | 1.2 | 90.0 | 1 |
| | 3180 | | 2 | 50.0 | 110.0 | 0.5 | 4.5455 | 0.1 | 2.4 | 1.2 | 100.0 | 2 |
| | 3929 | | 3 | 50.0 | 110.0 | 0.5 | 4.5455 | 0.1 | 2.4 | 1.8 | 100.0 | 3 |
| | 4523 | | 4 | 50.0 | 110.0 | 0.5 | 4.5455 | 0.1 | 2.4 | 2.4 | 100.0 | 4 |
| | 5599 | | 5 | 50.0 | 110.0 | 0.5 | 4.5455 | 0.1 | 2.4 | 3.0 | 100.0 | 6 |
| | 5613 | | 6 | 50.0 | 100.0 | 0.5 | 5.0 | 0.1 | 1.6 | 3.0 | 100.0 | 5 |
| | 6068 | | 7 | 50.0 | 10.0 | 0.1 | 5.0 | 0.1 | 2.4 | 3.0 | 80.0 | 7 |
| | 6177 | | 8 | 50.0 | 100.0 | 0.5 | 5.0 | 0.1 | 2.4 | 3.0 | 100.0 | 10 |
| | 6778 | | 9 | 50.0 | 100.0 | 0.5 | 5.0 | 0.1 | 2.4 | 3.0 | 90.0 | 9 |
| | 6785 | | 10 | 0.0 | 100.0 | 0.5 | 5.0 | 0.1 | 3.2 | 3.0 | 100.0 | 14 |
| | 6789 | | 11 | 0.0 | 110.0 | 0.5 | 4.5455 | 0.1 | 2.4 | 1.8 | 100.0 | 8 |
| (*racE*-) to (agg) comparison | | Intermediate layer | | - | n/a | - | n/a | ** | *** | n/a | ** | |
| (*racE*-) to (veg) comparison | | Intermediate layer | | - | n/a | - | n/a | * | - | n/a | ** | |

*… (p < 10%), **… (p < 0.1%), ***… (p < 0.001%), -… not significant
n/a : subject to bias from the choice of parameter search

**Table S10. Parameters for representative data in main figures (Fig. 2, 3 and 5)**

| Description | Parameters |
|---|---|
| Fig. 2b | $\chi_U = 50$, $\gamma = 0.1$, $\mu = 0.5$, $D_W = 3$, $\rho = 4.55$, $k_{W1} = 110$, $W_{tot} = 80$, $a_W = 2.4$, |
| Fig. 2c | $\chi_U = 50$, $\gamma = 0.1$, $\mu = 0.7$, $D_W = 3$, $\rho = 4.76$, $k_{W1} = 105$, $W_{tot} = 80$, $a_W = 2.4$ |
| Fig. 2d | $\chi_U = 50$, $\gamma = 0.1$, $\mu = 0.5$, $D_W = 3$, $\rho = 5.56$, $k_{W1} = 90$, $W_{tot} = [60, 65, 70, 75, 80, 85, 90, 95, 100, 105, 110]$, $a_W = 2.4$ |
| Fig. 2e (black) | $\chi_U = 50$, $\gamma = 0.1$, $\mu = 0.5$, $D_W = 3$, $\rho = 5.56$, $k_{W1} = 90$, $W_{tot} = [60, 65, 70, 75, 80, 85, 90, 95, 100, 105, 110, 115]$, $a_W = 2.4$ |
| (blue) | $\chi_U = 50$, $\gamma = 0.1$, $\mu = 0.1$, $D_W = 3$, $\rho = 5$, $k_{W1} = 100$, $W_{tot} = [50, 55, 60, 65, 70, 75, 80, 85, 90, 95, 100, 105, 110, 115]$, $a_W = 2.4$ |
| (magenta) | $\chi_U = 50$, $\gamma = 0.1$, $\mu = 0.5$, $D_W = 3$, $\rho = 4.55$, $k_{W1} = 110$, $W_{tot} = [65, 70, 75, 80, 85, 90, 95, 100, 105, 110, 115, 120, 125, 130]$, $a_W = 4$ |
| Fig. 2f | $\chi_U = 30$, $\gamma = 0.1$, $\mu = 0.5$, $D_W = 3$, $\rho = 4.55$, $k_{W1} = 110$, $W_{tot} = 80$, $a_W = [0.8, 1.6, 2.4, 3.2, 4]$ |
| Fig. 2g (black) | $\chi_U = 30$, $\gamma = 0.1$, $\mu = 0.5$, $D_W = 3$, $\rho = 4.55$, $k_{W1} = 110$, $W_{tot} = 80$, $a_W = [0.8, 1.6, 2.4, 3.2, 4, 4.8, 5.6]$ |
| (blue) | $\chi_U = 50$, $\gamma = 0.1$, $\mu = 0.1$, $D_W = 3$, $\rho = 5$, $k_{W1} = 100$, $W_{tot} = 80$, $a_W = [0.8, 1.6, 2.4, 3.2, 4, 4.8, 5.6]$ |
| Fig. 2h | $\chi_U = 50$, $\gamma = 0.1$, $\mu = 0.5$, $D_W = 3$, $\rho = 4.55$, $k_{W1} = [1.1, 2.2, 11, 55, 110]$, $W_{tot} = 80$, $a_W = 5.6$ |
| Fig. 2i (black) | $\chi_U = 50$, $\gamma = 0.1$, $\mu = 0.5$, $D_W = 3$, $\rho = 4.55$, $k_{W1} = [1.1, 2.2, 11, 22, 55, 110]$, $W_{tot} = 80$ $a_W = 5.6$ |
| (blue) | $\chi_U = 50$, $\gamma = 0.1$, $\mu = 0.5$, $D_W = 3$, $\rho = 4.55$, $k_{W1} = [1.1, 2.2, 11, 22, 55, 110]$, $W_{tot} = 90$ $a_W = 2.4$ |
| (magenta) | $\chi_U = 50$, $\gamma = 0.1$, $\mu = 0.1$, $D_W = 3$, $\rho = 5$, $k_{W1} = [1, 2, 10, 20, 50, 100]$, $W_{tot} = 80$, $a_W = 2.4$ |
| Fig. 2j | $\chi_U = 50$, $\gamma = 0.1$, $\mu = [0.1, 0.3, 0.5, 0.7, 0.9]$, $D_W = 3$, $\rho = 5$, $k_{W1} = 100$, $W_{tot} = 80$, $a_W = 2.4$ |
| Fig. 2k (black) | $\chi_U = 50$, $\gamma = 0.1$, $\mu = [0.1, 0.3, 0.5, 0.7, 0.9]$, $D_W = 3$, $\rho = 5$, $k_{W1} = 100$, $W_{tot} = 80$, $a_W = 2.4$ |
| (blue) | $\chi_U = 50$, $\gamma = 0.1$, $\mu = [0.02, 0.05, 0.1, 0.3, 0.5, 0.7, 0.9]$, $D_W = 3$, $\rho = 4.55$, $k_{W1} = 110$, $W_{tot} = 80$, $a_W = 4.8$ |
| Fig. 3c-e | $\chi_U = 0$, $\gamma = 0.3$, $\mu = 0.5$, $D_W = 1.2$, $\rho = 4.55$, $k_{W1} = 110$, $W_{tot} = 80$, $a_W = 4$ |
| Fig. 3f-h | $\chi_U = 50$, $\gamma = 0.1$, $\mu = 0.5$, $D_W = 3$, $\rho = 5$, $k_{W1} = 100$, $W_{tot} = 80$, $a_W = 1.6$ |
| Fig. 3i-k | $\chi_U = 50$, $\gamma = 0.1$, $\mu = 0.1$, $D_W = 3$, $\rho = 5$, $k_{W1} = 20$, $W_{tot} = 80$, $a_W = 2.4$ |
| Fig. 5c | $\chi_U = 0$, $\gamma = 0.1$, $\mu = 0.9$, $D_W = 3$, $\rho = 5$, $k_{W1} = 50$, $W_{tot} = 80$, $a_W = 2.4$ |
| Fig. 5e | $\chi_U = 50$, $\gamma = 0.1$, $\mu = 0.5$, $D_W = 1.2$, $\rho = 4.55$, $k_{W1} = 110$, $W_{tot} = 90$, $a_W = 2.4$ |
| Fig. 5g | $\chi_U = 0$, $\gamma = 0.1$, $\mu = 0.5$, $D_W = 3$, $\rho = 5$, $k_{W1} = 100$, $W_{tot} = 90$, $a_W = 1.6$ |

**Table S11 Parameters for representative data in supplementary figures (fig. S3, S4, S5 and S6)**

| Description | Parameters |
|---|---|
| fig. S3b | $\alpha = 10, \beta = 3.5, \chi_U = 50, \gamma = 0.32, \mu = 1.0, \theta = 0.014, d = 0.8, \sigma = 7.5, a = 35$, b=0.09 |
| fig. S3c | $\alpha = 10, \beta = 3.5, \chi_U = 50, \gamma = 0.32, \mu = 1.0, \theta = 14, d = 0.8, \sigma = 0.02, a = 35$, b=0.09 |
| fig. S3f | $D_W = 3, \rho = 5.56, k_{W1} = 90, W_{tot} = 80, \theta = 2.0, d = 5.0, \sigma = 0.75, a_W = 2.4$ |
| fig. S3g | $D_W = 3, \rho = 6.25, k_{W1} = 80, W_{tot} = 90, \theta = 1.4, d = 0.8, \sigma = 0.15, a_W = 0.8$ |
| fig. S4b R1 | $\alpha = 2, \beta = 3.5, \chi_U = 0, \gamma = 0.1, \mu = 0.5, D_W = 1.2, \rho = 4.55, k_{W1} = 110, W_{tot} = 80, a_W = 2.4$ |
| fig. S4b (1) | $\alpha = 2, \beta = 3.5, \chi_U = 0, \gamma = 0.5, \mu = 0.5, D_W = 1.2, \rho = 4.55, k_{W1} = 110, W_{tot} = 80, a_W = 2.4$ |
| fig. S4b (2) | $\alpha = 2, \beta = 3.5, \chi_U = 0, \gamma = 0.7, \mu = 0.5, D_W = 1.2, \rho = 4.55, k_{W1} = 110, W_{tot} = 80, a_W = 2.4$ |
| fig. S4b (3) | $\alpha = 2, \beta = 3.5, \chi_U = 0, \gamma = 0.5, \mu = 0.5, D_W = 1.2, \rho = 4.55, k_{W1} = 110, W_{tot} = 80, a_W = 0.8$ |
| fig. S4b (4) | $\alpha = 2, \beta = 3.5, \chi_U = 0, \gamma = 0.5, \mu = 0.5, D_W = 1.2, \rho = 4.55, k_{W1} = 110, W_{tot} = 80, a_W = 4$ |
| fig. S4c R2' | $\alpha = 2, \beta = 3.5, \chi_U = 50, \gamma = 0.1, \mu = 0.5, D_W = 3, \rho = 5, k_{W1} = 100, W_{tot} = 80, a_W = 2.4$ |
| fig. S4c (5) | $\alpha = 2, \beta = 3.5, \chi_U = 50, \gamma = 0.1, \mu = 0.5, D_W = 3, \rho = 50, k_{W1} = 10, W_{tot} = 80, a_W = 2.4$ |
| fig. S4c (6) | $\alpha = 2, \beta = 3.5, \chi_U = 50, \gamma = 0.1, \mu = 0.5, D_W = 3, \rho = 2.5, k_{W1} = 200, W_{tot} = 80, a_W = 2.4$ |
| fig. S4c (7) | $\alpha = 2, \beta = 3.5, \chi_U = 50, \gamma = 0.1, \mu = 0.1, D_W = 3, \rho = 5, k_{W1} = 100, W_{tot} = 80, a_W = 2.4$ |
| fig. S4c (8) | $\alpha = 2, \beta = 3.5, \chi_U = 50, \gamma = 0.1, \mu = 0.9, D_W = 3, \rho = 5, k_{W1} = 100, W_{tot} = 80, a_W = 2.4$ |
| fig. S4d | $\alpha = 2, \beta = 3.5, \chi_U = 0, \gamma = [0.1, 0.3, 0.5, 0.7], \mu = 0.5, D_W = 1.2, \rho = 4.55, k_{W1} = 110, W_{tot} = 80, a_W = [0.8, 1.6, 2.4, 3.2, 4]$ |
| fig. S4e | $\alpha = 2, \beta = 3.5, \chi_U = 50, \gamma = 0.1, \mu = [0.1, 0.3, 0.5, 0.7, 0.9], D_W = 3, \rho = 5, k_{W1} = [10, 20, 50, 100, 200], W_{tot} = 80, a_W = 2.4$ |
| fig. S5a, b | $\alpha = 2, \beta = 3.5, \chi_U = 50, \gamma = 0.1, \mu = 0.5, D_W = 3, \rho = [4.55, 4.76, 5, 5.26, 5.56], k_{W1} = 500, W_{tot} = 80, a_W = 2.4$ |
| fig. S5c, d | $\alpha = 2, \beta = 3.5, \chi_U = 0, \gamma = [0.1, 0.3, 0.5, 0.7, 0.9], \mu = 0.5, D_W = 3, \rho = 4.55, k_{W1} = 110, W_{tot} = 80, a_W = 2.4$ |
| fig. S5e, f | $\alpha = 2, \beta = 3.5, \chi_U = 0, \gamma = 0.1, \mu = 0.5, D_W = [0.6, 1.2, 1.8, 2.4, 3, 3.6], \rho = 4.55, k_{W1} = 110, W_{tot} = 80, a_W = 2.4$ |
| fig. S5g, h | $\alpha = 2, \beta = 3.5, \chi_U = [0, 10, 30, 50], \gamma = 0.1, \mu = 0.5, D_W = 3, \rho = 4.55, k_{W1} = 110, W_{tot} = 80, a_W = 2.4$ |
| fig. S6c | $\alpha = 2, \beta = 3.5, \chi_U = 0, \gamma = 0.7, \mu = 0.5, D_W = 1.2, \rho = 4.55, k_{W1} = 110, W_{tot} = 80, a_W = 4$ |
| fig. S6d | $\alpha = 2, \beta = 3.5, \chi_U = 0, \gamma = 0.5, \mu = 0.5, D_W = 3, \rho = 5.56, k_{W1} = 90, W_{tot} = 80, a_W = 2.4$ |

**Supplementary Figure legends:**

**Figure S1. Image classification of migratory cells based on a deep convolutional neural network provides highly compressed representation of the overall cell shape, protrusions and their orientation.** (a, b) The values of accuracy (a) and loss (b) during training (blue) and validation (red) of the deep convolutional neural networks. (c-e) Representative time series of the feature vector $\boldsymbol{F}$ for *Dictyostelium* (agg)(c), HL60 (d) and fish keratocyte (e). (f, g) Orientation dependency for the oval shape (Fig 1(m)) $L = 0.75L_0$ (f) and $L = L_0$ (g). (h-j) Mapping of circles and polygons with various aspect ratios (x-axis: y-axis) 1:3, 2:5, 1:2, 2:3, 1:1, 3:2, 2:1, 5:2 and 3:1. Circles, squares and rhombuses (h), triangles, pentagons and star shapes in the upright (i) and inverted (j) orientation. (k) Orientation dependency of a complex shape with multiple edges.

**Figure S2. Analysis of real cell data with varying degree of bifurcating protrusions indicates polar and multiple edge representation in PC2.** (a, b) Mapping of cell shape with anterior-posterior elongation without lateral pseudopods (*Dictyostelium* prespore cell-type). Cell-to-cell variation (a) and a representative temporal variation of a single cell (b). Blue circled regions represent 95% confidence eclipses for the mean of all combined timeseries (dotted) and the mean of individual cells (filled). (c, d) Mapping of mouse T cells in the PC1-PC2 space. Filled rhombuses: T helper 1 (c) and regulatory T cell (d). Circled regions in the background indicate the reference data in Fig. 1b (aggregation-stage *Dictyostelium* dark green (dark green) , HL60 (dark red) and fish keratocyte (yellow)).

**Figure S3. Model behaviors in the 2-variable and 1-variable limit shows other repertoires in the morphodynamics.** : (a-d) The 2-variable scheme (a) and its morphology dynamics (b-d). Representative simulations showing traveling patches (b) and lamellipodium-like protrusions (c). Color overlay; red $V$, green $U$. (d) Time series of the local curvature of the boundary (left panel) and local elongation (right panel) for (c). (e-g) The 1-variable scheme (e) and its representative morphology dynamics (f, g). Blue indicates the high $W$ region. (h) Time series of the local curvature at the boundary (let panel) and local elongation (right panel) for (g). (i) Feature mapping of the polar morphology in 1-variable scheme. Time average (solid circles) and time samples (filled). Right panels in colored frames indicate representative snapshots. See Table S11 for parameter values.

**Figure S4. Sampled parameter conditions.** (a) Two-dimensional grids around manually selected reference parameters R1 (upper left panels; $\chi_U = 0$), R1' (lower left panels, $\chi_U = 50$) , R2 (upper right panels) and R2' (lower right panels). Each square in the grid represents a sampled parameter condition. (b, c) Representative simulation time series for parameters R1, R2' and (1)-(8) in (a). (d) Feature scores ($F_1$, $F_2$, $F_3$) in the ($\gamma$-$a_W$ ) plane around R1. (e) Feature scores ($F_1$, $F_2$, $F_3$) in the ($k_{W1}$-$\mu$) plane around R2'. See Table S11 for parameter values.

**Figure S5. Parameter dependency of the simulated morphology**: Dependency on $\gamma$, $\rho$, $D_W$ , $\chi_U$ of the simulated morphology (a, c, e, g) and the feature values (b, d, f, h). See Table S11 for parameter values.

**Figure S6 An intermediate-layer of the convolutional neural network yields a similar feature space.** (a) PCA of the 256 dimensional intermediate layer. Microscopy dataset of *Dictyostelium* (agg) (green +), HL-60 (dark red +) , keratocyte (yellow +), *Dictyostelium* (veg) (magenta inverted triangles), *Dicytostelium racE*- strain (red squares) and Nocodazole-treated HL-60 (blue triangles). (b) Mapping of model simulations (grey circles), the top-ranking simulations based on the feature vector $\boldsymbol{F}$ (dark green circle for *Dictyostelium* (agg), magenta circle for *Dictyostelium* (veg)) and those based on the 256-dimensional features (light green circle for *Dictyostelium* (agg), pink circle for *Dictyostelium* (veg)). (c,d) Time-series of simulations with the highest feature similarity to *Dictyostelium* (aggregation-stage) (c), vegetative *Dicyostelium* (d). See Table S11 for parameter values.

**Supplementary Movie legends:**

Movie S1: Timelapse sequence of representative training data images; *Dictyostelium* (agg), HL60 and fish keratocyte (from left to right).

Movie S2: Parameter dependency ($W_{\mathrm{tot}}$) of the morphodynamic model (Fig. 2d).

Movie S3: Parameter dependency ($k_{\mathrm{w1}}$) of the morphodynamic model (Fig. 2f).

Movie S4: Parameter dependency ($a_{\mathrm{w}}$) of the morphodynamic model (Fig. 2h).

Movie S5: Parameter dependency ($\mu$) of the morphodynamic model (Fig. 2j).

Movie S6: Cell morphology dynamics of aggregation stage *Dictyostelium* and the corresponding top ranking simulations.

Movie S7: Cell morphology dynamics of HL60 and the corresponding top ranking simulations.

Movie S8: Cell morphology dynamics of fish keratocyte and the corresponding top ranking simulations.

Movie S9: Cell morphology dynamics of vegetative stage *Dictyostelium* and the corresponding top ranking simulations.

Movie S10: Cell morphology dynamics of nocodazole-treated HL60 and the corresponding top ranking simulations.

Movie S11: Cell morphology dynamics of *Dictyostelium* racE- and the corresponding top ranking simulations.

Fig.S1

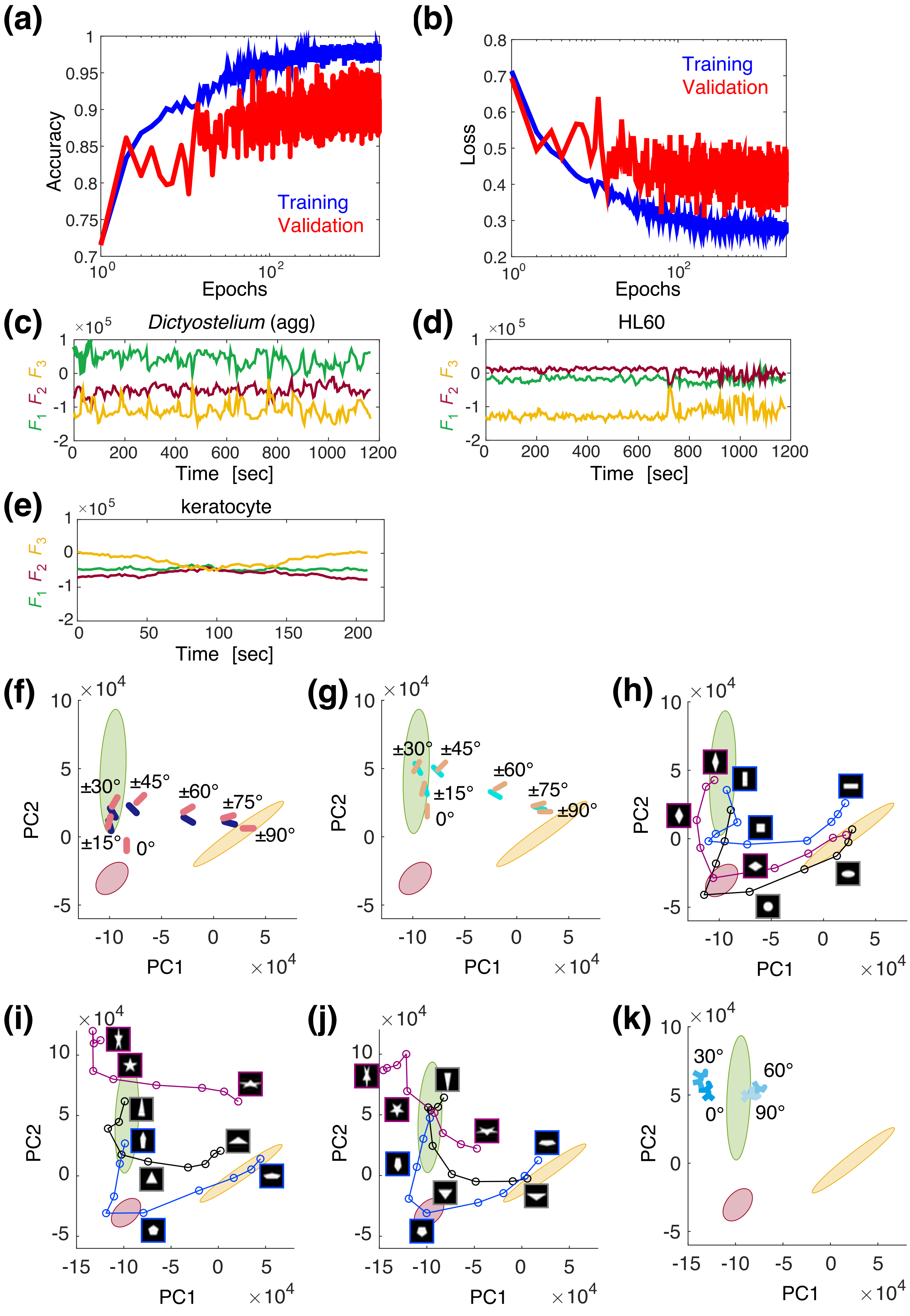

(a)
Accuracy
Epochs
Training
Validation
(b)
Loss
Epochs
Training
Validation
(c)
Dictyostelium (agg)
$F_1$ $F_2$ $F_3$
Time [sec]
(d)
HL60
$F_1$ $F_2$ $F_3$
Time [sec]
(e)
keratocyte
$F_1$ $F_2$ $F_3$
Time [sec]
(f)
PC1
PC2
±30° ±45° ±60° ±75° ±90° ±15° 0°
(g)
PC1
PC2
±30° ±45° ±60° ±15° ±75° ±90° 0°
(h)
PC1
PC2
(i)
PC1
PC2
(j)
PC1
PC2
(k)
PC1
PC2
30° 60° 0° 90°

Fig.S2

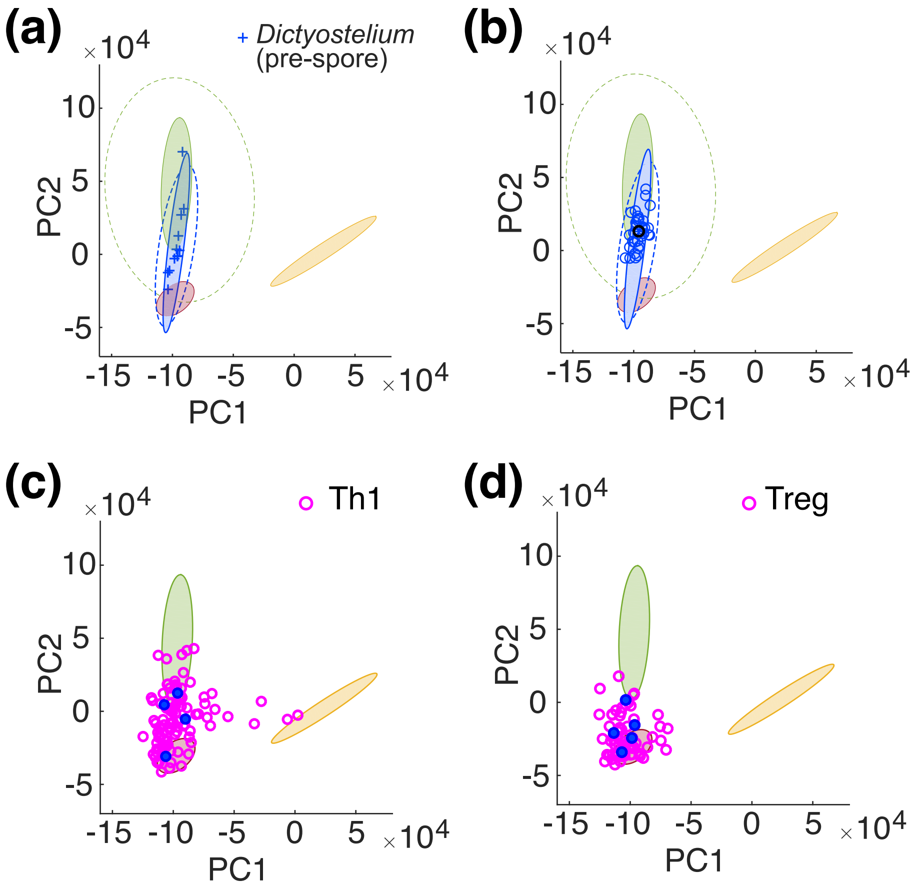

(a)
Dictyostelium
(pre-spore)
PC2
PC1
(b)
PC2
PC1
(c)
Th1
PC2
PC1
(d)
Treg
PC2
PC1

Fig.S3

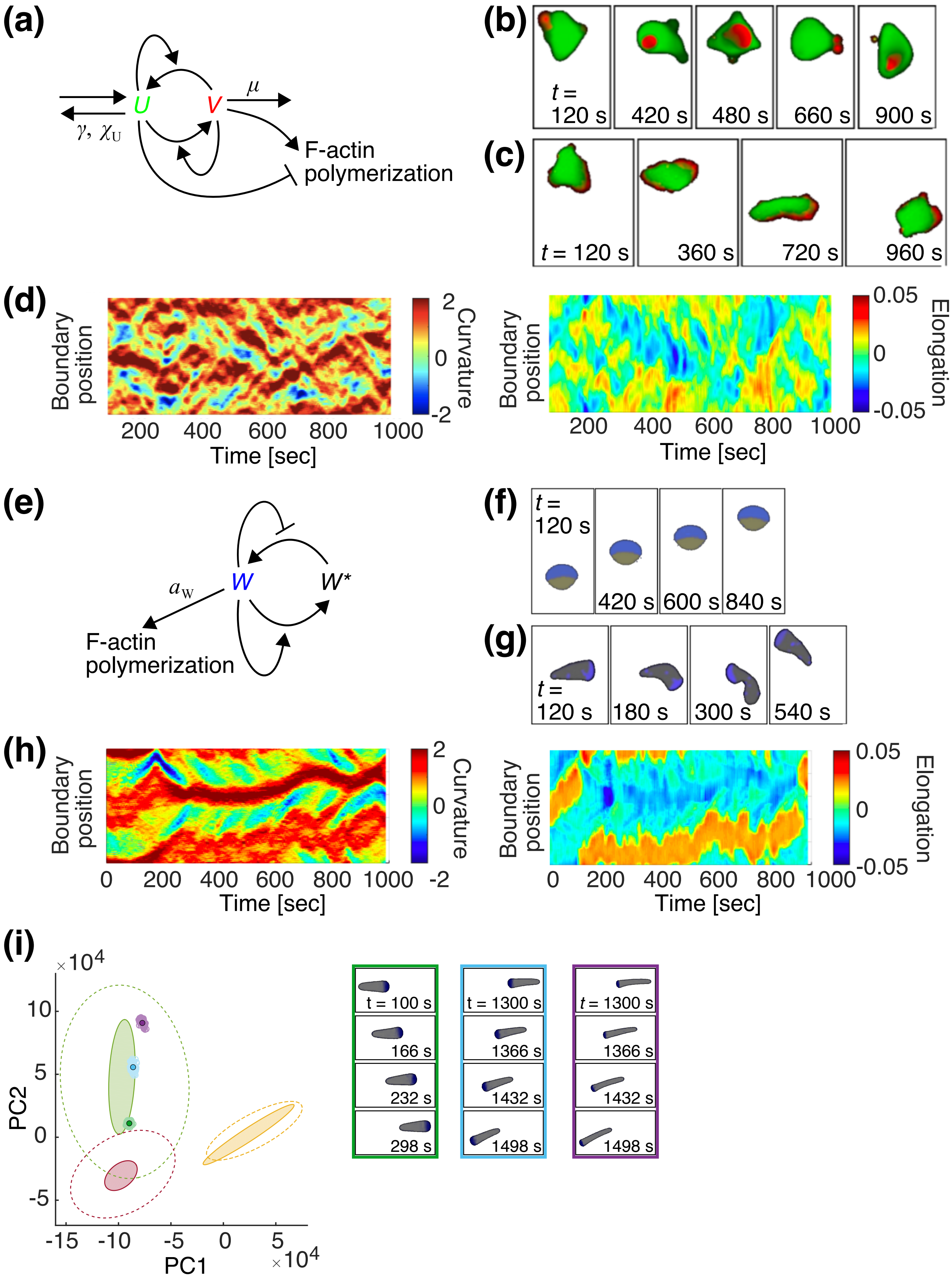

(a)
U
V
μ
γ, χU
F-actin
polymerization
(b)
t =
120 s
420 s
480 s
660 s
900 s
(c)
t = 120 s
360 s
720 s
960 s
(d)
Boundary position
Time [sec]
Curvature
Elongation
(e)
W
W*
aW
F-actin
polymerization
(f)
t =
120 s
420 s
600 s
840 s
(g)
t =
120 s
180 s
300 s
540 s
(h)
Boundary position
Time [sec]
Curvature
Elongation
(i)
PC1
PC2
×10^4
t = 100 s
166 s
232 s
298 s
t = 1300 s
1366 s
1432 s
1498 s
t = 1300 s
1366 s
1432 s
1498 s

Fig.S4

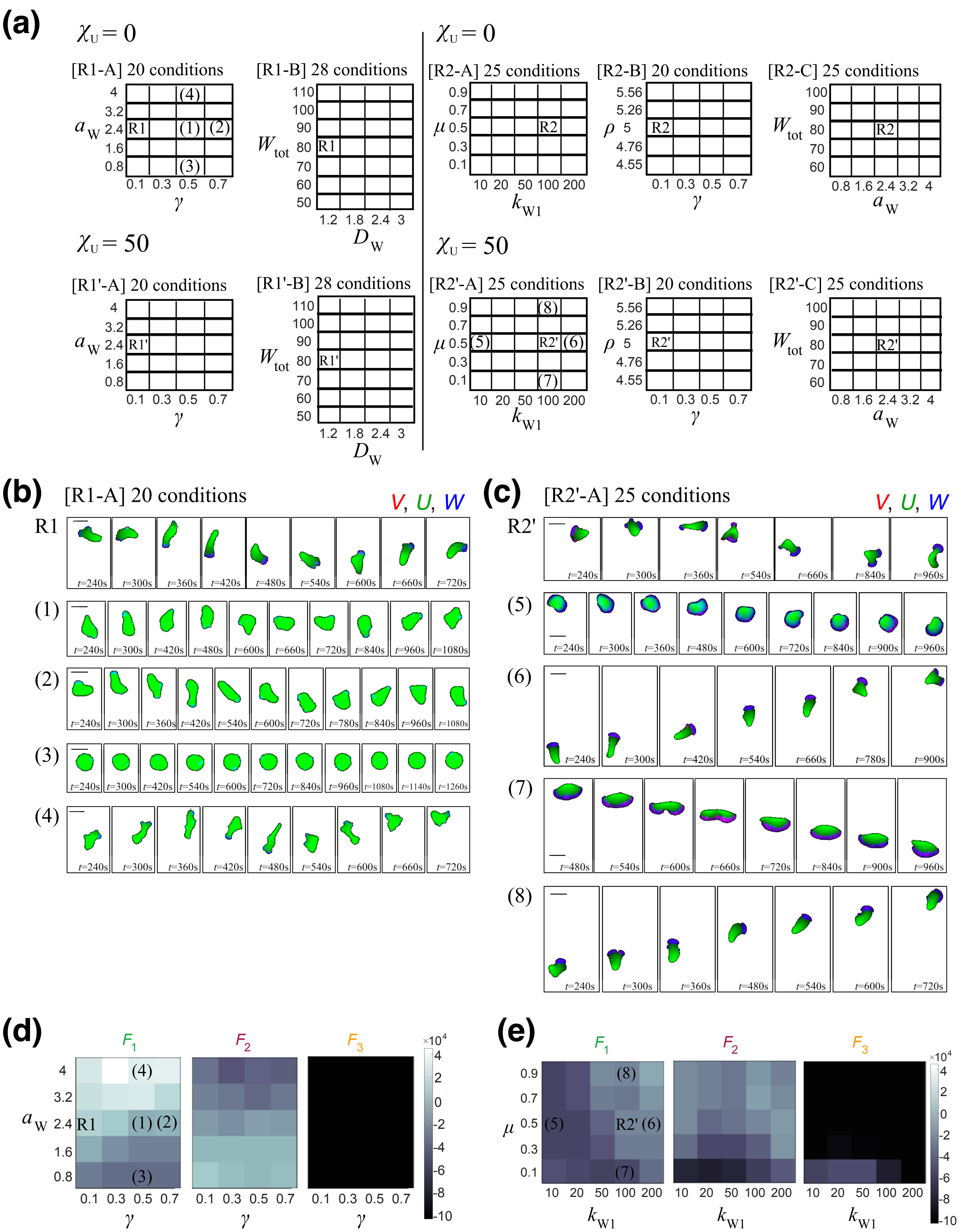
(a)
$\chi_U = 0$
[R1-A] 20 conditions
[R1-B] 28 conditions
[R2-A] 25 conditions
[R2-B] 20 conditions
[R2-C] 25 conditions
$\chi_U = 50$
[R1'-A] 20 conditions
[R1'-B] 28 conditions
[R2'-A] 25 conditions
[R2'-B] 20 conditions
[R2'-C] 25 conditions
(b)
[R1-A] 20 conditions
V, U, W
(c)
[R2'-A] 25 conditions
V, U, W
(d)
F1 F2 F3
(e)
F1 F2 F3

Fig.S5

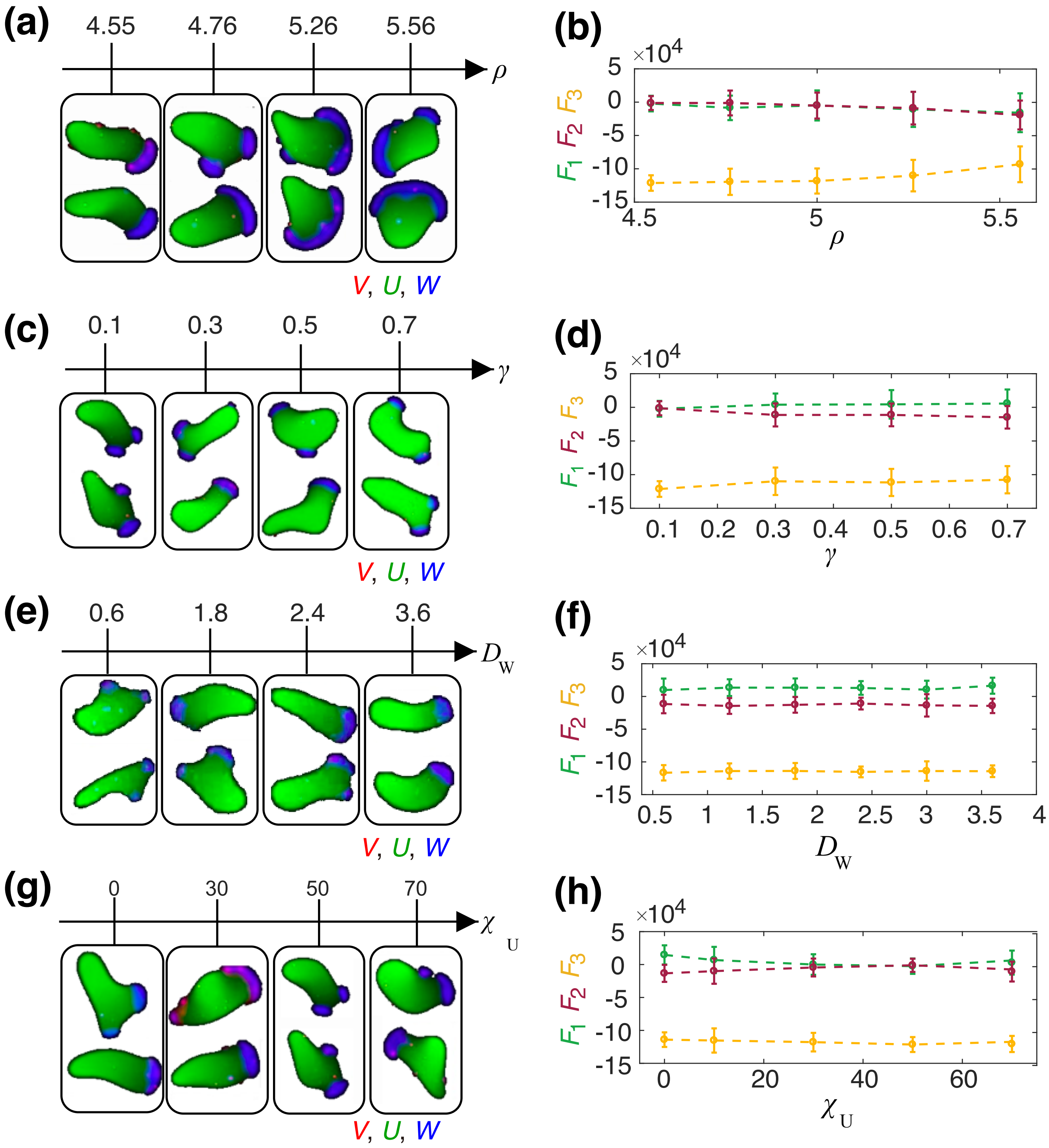
(a)
4.55
4.76
5.26
5.56
ρ
V, U, W
(b)
×10^4
5
0
-5
-10
-15
F1 F2 F3
4.5
5
5.5
ρ
(c)
0.1
0.3
0.5
0.7
γ
V, U, W
(d)
×10^4
5
0
-5
-10
-15
F1 F2 F3
0.1
0.2
0.3
0.4
0.5
0.6
0.7
γ
(e)
0.6
1.8
2.4
3.6
DW
V, U, W
(f)
×10^4
5
0
-5
-10
-15
F1 F2 F3
0.5
1
1.5
2
2.5
3
3.5
4
DW
(g)
0
30
50
70
χU
V, U, W
(h)
×10^4
5
0
-5
-10
-15
F1 F2 F3
0
20
40
60
χU

Fig.S6

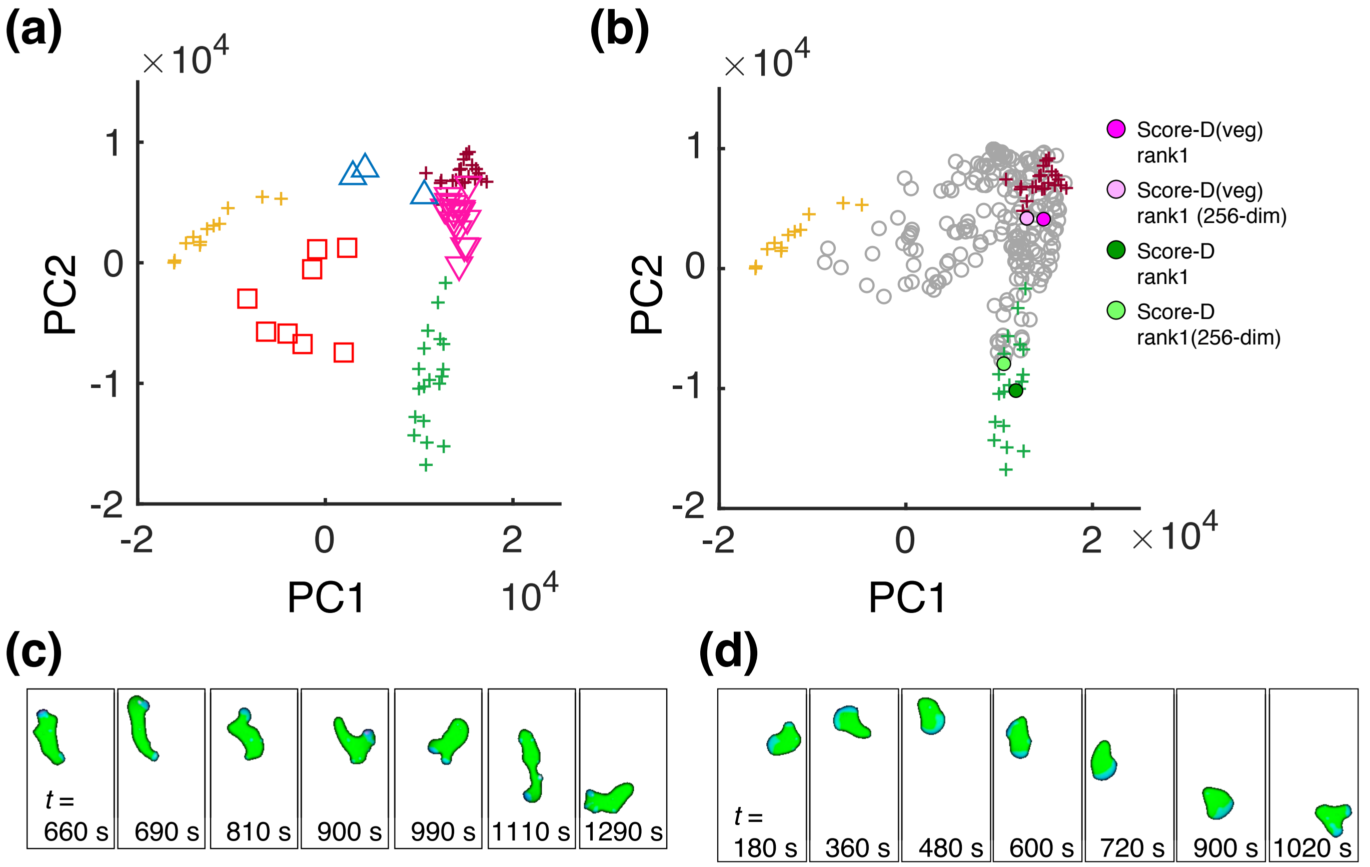
(a)
(b)
$\times 10^4$
PC2
PC1
1
0
-1
-2
2
$10^4$
Score-D(veg) rank1
Score-D(veg) rank1 (256-dim)
Score-D rank1
Score-D rank1(256-dim)
(c)
(d)
t =
660 s
690 s
810 s
900 s
990 s
1110 s
1290 s
180 s
360 s
480 s
600 s
720 s
900 s
1020 s
